# Differential localization patterns of pyruvate kinase isoforms in murine naïve, formative and primed pluripotent states

**DOI:** 10.1101/2020.04.12.036251

**Authors:** Joshua G. Dierolf, Andrew J. Watson, Dean H. Betts

**Affiliations:** Departments of Physiology and Pharmacology, The University of Western Ontario, London, Canada; Obstetrics and Gynecology, Schulich School of Medicine & Dentistry, The University of Western Ontario, London, Canada; The Children’s Health Research Institute (CHRI), Lawson Health Research Institute, London, Canada

## Abstract

Mouse embryonic stem cells (mESCs) and mouse epiblast stem cells (mEpiSCs) represent opposite ends of a pluripotency continuum, referred to as naïve and primed pluripotent states, respectively. These divergent pluripotent states differ in several ways including growth factor requirements, transcription factor expression, DNA methylation patterns, and metabolic profiles. Naïve cells employ both glycolysis and oxidative phosphorylation (OXPHOS), whereas primed cells preferentially utilize aerobic glycolysis, a trait shared with cancer cells referred to as the Warburg effect. Until recently, metabolism has been regarded as a by-product of cell fate; however, evidence now supports metabolism as being a driver of stem cell state and fate decisions. Pyruvate kinase muscle isoforms (PKM1 and PKM2) are important for generating and maintaining pluripotent stem cells (PSCs) and mediating the Warburg effect. Both isoforms catalyze the last step of glycolysis generating adenosine triphosphate and pyruvate, however, the precise role(s) of PKM1/2 in naïve and primed pluripotency is not well understood. The primary objective was to characterize the cellular expression and localization patterns of PKM1 and PKM2 in mESCs, chemically transitioned epiblast-like cells (mEpiLCs) representing formative pluripotency, and mEpiSCs using immunoblotting, flow cytometry and confocal microscopy. The results indicate that PKM1 and PKM2 are not only localized to the cytoplasm but also accumulate in differential subnuclear regions of mESC, mEpiLCs and mEpiSCs as determined by a quantitative, confocal microscopy colocalization employing orthogonal projections and airyscan processing. Importantly, we discovered that the subnuclear localization of PKM1/2 shifts during the transition from mESCs, mEpiLCs and mEpiSCs. We have also authenticated a new method of selecting formative pluripotency cells from naïve and primed populations using the cell surface markers SSEA1 and CD24. Finally, we have comprehensively validated the appropriateness and power of the Pearson’s correlation coefficient and Manders’ overlap coefficient for assessing nuclear and cytoplasmic protein colocalization in PSCs by immunofluorescence confocal microscopy. We propose that nuclear PKM1/2 assists with distinct pluripotency state maintenance and lineage priming by non-canonical mechanisms. These results advance our understanding of the overall mechanisms controlling naïve, formative and primed pluripotency.

## Introduction

Pluripotent stem cells (PSCs) have the capacity for indefinite self-renewal and the potential to differentiate into the cell types of all three germ layers including the germ line (1). The potency of PSCs, such as embryonic stem cells (ESCs), exists within a continuum with opposite ends described as naïve and primed states (1). In mice, naïve mESCs are derived from the inner cell mass (ICM) of an early, embryonic day (E)3.5 to 4.5, blastocyst-stage embryo, whereas primed mouse epiblast stem cells (mEpiSCs) are derived later from the epiblast of E5.0-8.0 post-implantation embryos (2–6). However, when cultured *in vitro*, mEpiSCs more closely resemble the epiblast of E7.25-8.0 embryos (2,5,7,8). Human ESCs (hESCs) have traditionally been stabilized at the primed pluripotent state, however, a naïve hESC line has been recently derived (9). Between both ends of the pluripotent continuum exists a recently described intermediate state called the “formative pluripotent state” (10,11). Formative pluripotency is an executive phase and may represent the gene expression patterns and attributes of mouse epiblast cells within E5.5-6.25 embryo (2). Like naïve and primed PSCs, formative PSCs also express NANOG, OCT4 and SOX2 (10,12,13). However, unlike naïve and primed PSCs, the formative mouse epiblast-like cells (mEpiLCs) can efficiently differentiate into primordial germ cell-like cells when presented with the appropriate growth factors such as bone morphogenic protein 4 (2,14). Each pluripotent state has several distinguishing features such as unique morphology, growth factor dependencies, gene expression profiles, epigenetic status and metabolic preferences (1,2). Morphologically, naïve PSCs are more rounded in appearance and grow as colonies with glistening edges compared to flattened primed PSC colonies (1). This hemispherical morphology of naïve cells is largely due to greater Cdh1 expression, which can be replicated in mEpiSCs following overexpression of Cdh1 (15). Culture of mESCs requires leukemia inhibitor factor (LIF) which promotes ‘ground state’ naïve pluripotency and resists differentiation through activation of the transcription factor STAT3 (16). Stabilizing naïve pluripotency requires LIF and the addition of two small molecule inhibitors (LIF/2i) of MEK1/2 (PD00032) and glycogen synthase kinase-3 (CHIR99021) (17,18). Formative cells can be chemically transitioned from mESCs to mouse epiblast-like cells (mEpiLCs) over 48 hours as a transient and heterogenous population (13,19). To maintain primed pluripotency and exit the naïve state, mEpiSCs and chemically transitioned mEpiLCs are cultured with ACTIVIN-A and FGF-2 (3). Currently, there are no defined ways to induce and stably maintain the culture of the transient formative state mEpiLCs (19). While naïve and primed cells express the core pluripotency associated genes Oct4, Sox2, Nanog, both states differ in transcriptional programs with Rex1, Esrrb, Dppa3, Klf2/4/5, Tcfcp2l1 and Pecam delineating the naïve state, and Zic2, T (Brachyury) and Cer1, to list a few, distinguishing the primed pluripotent state (2). The formative pluripotent state is reported to highly express Lef1, Pou1fc and Dnmt3 (20). Naïve and primed pluripotent states also differ in terms of their epigenetic landscape, including X-activation and chromatin methylation status (21). Female primed PSCs display random X chromosome inactivation, whereas naïve PSCs display activation of both X chromosomes (22). Relative to primed cells, naïve PSCs contain larger regions of active chromatin as indicated by higher levels of H2k4me3 and histone acetylation (23,24). Importantly, naïve and primed PSCs also differ in terms of their metabolic preferences (25). Naïve cells are characterised as being metabolically bivalent, utilizing both glycolytic and oxidative phosphorylation (OXPHOS) processes, whereas primed cells are preferentially glycolytic (25). Even when cultured in oxygen rich conditions, primed PSCs utilize aerobic glycolysis and display low OXPHOS gene expression, which is characteristic of the Warburg effect that is activate in many cancer cells (26).

Cells exhibiting the Warburg effect not only consume glucose preferentially, but direct pyruvate towards lactate formation over a mitochondrial fate in OXPHOS, thus producing less carbon dioxide (27). These cells may not necessarily generate as much ATP through OXPHOS but generate intermediates that are used in biosynthesis, such as citrate, which is necessary to generate acetyl Co-A for downstream processes and non-essential amino acids (28). The Warburg effect is orchestrated by an upregulation of key transcription factors including: Oct4, c-Myc, Hif-1∝ and Nf*κ*b along with the glycolytic genes: Hk2, Pgm, Pdk and pyruvate kinase muscle isoform 2 (Pkm2) (29). When upregulated, these Warburgian drivers enhance anabolism and ATP generation to boost glycolytic flux (28,30). It is hypothesized that the high glycolytic flux of mESC maintains their high proliferative capacity and as such, cellular metabolic state should be considered as a mediator of pluripotency and as a regulator of gene expression controlling cell proliferation and differentiation (31,32). While metabolism has traditionally been viewed as a by-product of cell fate decisions, the manipulation of metabolic genes and their products in stem cells can promote or resist cellular differentiation and reprogramming (25,33,34). Thus, the developmental progression of naïve to primed transitioning occurs in synchrony with metabolic programming to influence cell fate and pluripotent state as both a driver and a passenger (35).

Recently, pyruvate kinase muscle isoforms 1 and 2 (PKM1/2) have been implicated in regulating pluripotency, proliferation and in the generation of pluripotent stem cells during reprogramming (36). PKM1 and PKM2 are the metabolic enzymes responsible for catalyzing the rate limiting step of glycolysis by directing pyruvate towards either a lactate or acetyl-CoA fate (37,38). Mammals express four tissue specific pyruvate kinase isozymes; M1, M2, L and R, each with unique properties and tissue expression patterns to meet specific metabolic demands (39). PKM1/2 are alternatively spliced isoforms from the PKM gene and both PKL and PKR are encoded by the PKLR gene (37). The M1 and M2 isoforms are spliced by three different heterogenous nuclear ribonucleoproteins; hnRNPI/hnRNP1/hnRNP2 that involve the inclusion of exon 9 or 10, respectively (40). PKM1/2 activity is regulated by homotropic and heterotropic allosteric interactions with fructose 1,6-bisphosphate (FBP) and phosphoenolpyruvate respectively (41,42). PKM1 is expressed primarily in somatic cells, whereas fetal tissues along with essentially all cancer cell types exhibit elevated PKM2 with certain types of tumours such as glioblastomas displaying a complete isoform switch from PKM1 to PKM2 (43). The elevated PKM2 found in cancer cells is predominantly the inactive PKM2 homodimer form, which is due to pulsatile phosphofructokinase (44). The active homotetramer is typically bound to its cofactor FBP, however, when the PKM2 homodimer is phosphorylated (Y105) by the oncogenic linked fibroblast growth factor receptor type 1, the homotetrameric configuration is disrupted (45,46). This interrupts glucose oxidation and increases glycolysis and lactate production in Warburgian cancer cells, even in the presence of abundant oxygen levels. In contrast, PKM1 operates as a constitutively active homotetramer without a described allosteric binding site (47).

PKM2 has additional non-canonical roles including its function as a protein kinase, cytosolic receptor, transcriptional co-activator and is also implicated in cytokinesis and chromosome segregation (48–50). PKM2 can form a complex with OCT4 resulting in decreased OCT4 transcriptional activity and stemness with increased apoptosis and differentiation (51,52). Studies also indicate that the interaction of PKM2 and OCT4 affects mitosis and tumor energy production (53). PKM2 is implicated in pluripotency through its regulation of OCT4 in hESCs (54). Knockdown of PKM2 in hESCs exhibited no change in lactate production or glucose uptake, however OCT4 expression decreased substantially (54). PKM2 is observed in the nuclei of the hESCs cultured under both normoxic (20%) and hypoxic (5%) oxygen conditions but a significant reduction in PKM2 expression was observed under normoxia (54). Overexpression of either PKM1 or PKM2 results in increased transcript abundance of the pluripotency associated genes; Eras, Rex1 and Nanog in mESCs (36). Upon knockdown of total PKM, pluripotency associated gene transcript abundance also decreases but self-renewal and morphology appear unperturbed (36). During reprogramming of somatic cells into iPSCs, both PKM1 and PKM2 are upregulated within the first 8 days (36). Additionally, the knockdown of total PKM during this period hinders reprogramming and overexpression of PKM2 significantly increases the generation of iPSC colonies (36). Interestingly, PKM1 has recently been localized in the nuclei of hepatoma (HepG2 and SMMC-7721 cell lines) cells following oroxylin A (OA) treatment and this localization was concluded to promote cellular differentiation to hepatocytes-like cells (55).

The mechanisms controlling PKM1 and PKM2 nuclear translocation are largely unknown, however, PKM1 may complex with hepatocyte nuclear factor 4 ∝ and this can be enhanced with the addition of the drug OA and the oncogene JMJD5 is implicated in the nuclear translocation of PKM2 (55,56). Nuclear translocation of PKM2 is well supported by fluorescent imaging and nuclear/cytoplasmic fractionation (57–61). However, typical confocal image analysis employing visual interpretation of overlaid fluorescent images is a purely qualitative means of spatial localization. Accurate quantitative measurement of spatial localization can effectively be quantified by a well-controlled comparison of two fluorophores to determine the degree of colocalization (62). Quantitative colocalization analysis (QCA) is most commonly divided into two metrics representing the relationship between two fluorophores, these measures are the degree of overlap and correlation (63). The degree of spatial location by overlapping images was first quantified by Otsu in 1979 where pixels of two images were overlapped after applying a threshold (64). Manders’ overlap coefficient (MOC) better distinguishes pixels ignored from the threshold from higher intensity pixels but at the cost of being influenced by autofluorescence and an insensitively to differences between the signal-to-noise ratios of the two fluorophores (65,66). While the MOC is a measure of co-occurrence of two fluorophores, within the spatially shared regions of a cell, two markers may interact or share a similar trend in intensity localization and may be functionally related or interact. Thus, the colocalization metric of correlation can indicate that two fluorophores share an associative relationship (66). The Pearson correlation coefficient (PCC) compares the variation of signal intensity between the intersection of two images taking into account the total population of pixels (66). As such, this calculation can determine the direction of linear association between the fluorophores (66,67).

Both MOC and the PCC are commonly used to quantify fluorescent protein spatial overlap and correlation (68). Despite the accuracy and power of QCA, this technique has been not been utilized to its full extent, especially so, in its application to measuring protein colocalization in pluripotent stem cells (63). This may be due to an on-going debate within the QCA field over the correct use and interpretation of overlap and correlation metrics (66,69,70). Thus, our study contrasted both PCC versus MOC in our analysis of PKM1/2 colocalization with nuclear and cytoplasmic protein markers during naïve, formative and primed pluripotency within mouse ES cell cultures (71).

We have, for the first time, comprehensively characterized the subcellular localization and expression patterns of PKM1 and PKM2 isoforms during transition from naïve, through the formative and onto the primed murine embryonic pluripotent states. We accomplished this characterization by optimizing a confocal microscopy colocalization approach comparing correlation and co-occurrence of PKM1 and PKM2 localization to nuclear localized OCT4 and cytoplasmic localized GAPDH. Degrees of colocalization were then applied to our measured values of overlap and correlation using qualified ranges indicating a spectrum of ‘very weak’ to ‘very strong’ variables of colocalization (71). Using these approaches, we report an elevated nuclear presence of PKM1 and PKM2 in naïve mESCs, formative state mEpiLCs and primed mEpiSCs as assessed by spatial overlap of PKM1 and PKM2 localization to OCT4 localization. We also report a moderate association of PKM1 and PKM2 to OCT4 localization in naïve mESCs and a strong association between PKM1 and OCT4 in formative mEpiLCs. Using nuclear and cytoplasmic fractionation, we verified increased protein abundance in the nuclear fraction of mESCs. Together, our results suggest a novel, non-canonical role for PKM1 in pluripotent stem cells.

## Results

### Characterization of naïve mESCs transitioning towards a primed pluripotent state

By 72 hours following the removal of mouse LIF and 2i supplementation with the addition of Fgf-2/Activin A (FA media), mESCs approximating a primed-like pluripotent state underwent an apoptotic event with the resulting colonies transitioning to a flattened morphology (S1 Fig). The mESCs by 72 hours had transitioned to mEpiLCs (primed-like state) and the mESCs, mEpiLCs and mEpiSCs showed homogenous colony expression of the pluripotency associated genes NANOG, OCT4 and SOX2 (Fig. 1A). Secondary antibody only controls confirmed the specificity of the immunofluorescence staining (S2 Fig).

**Fig 1.**
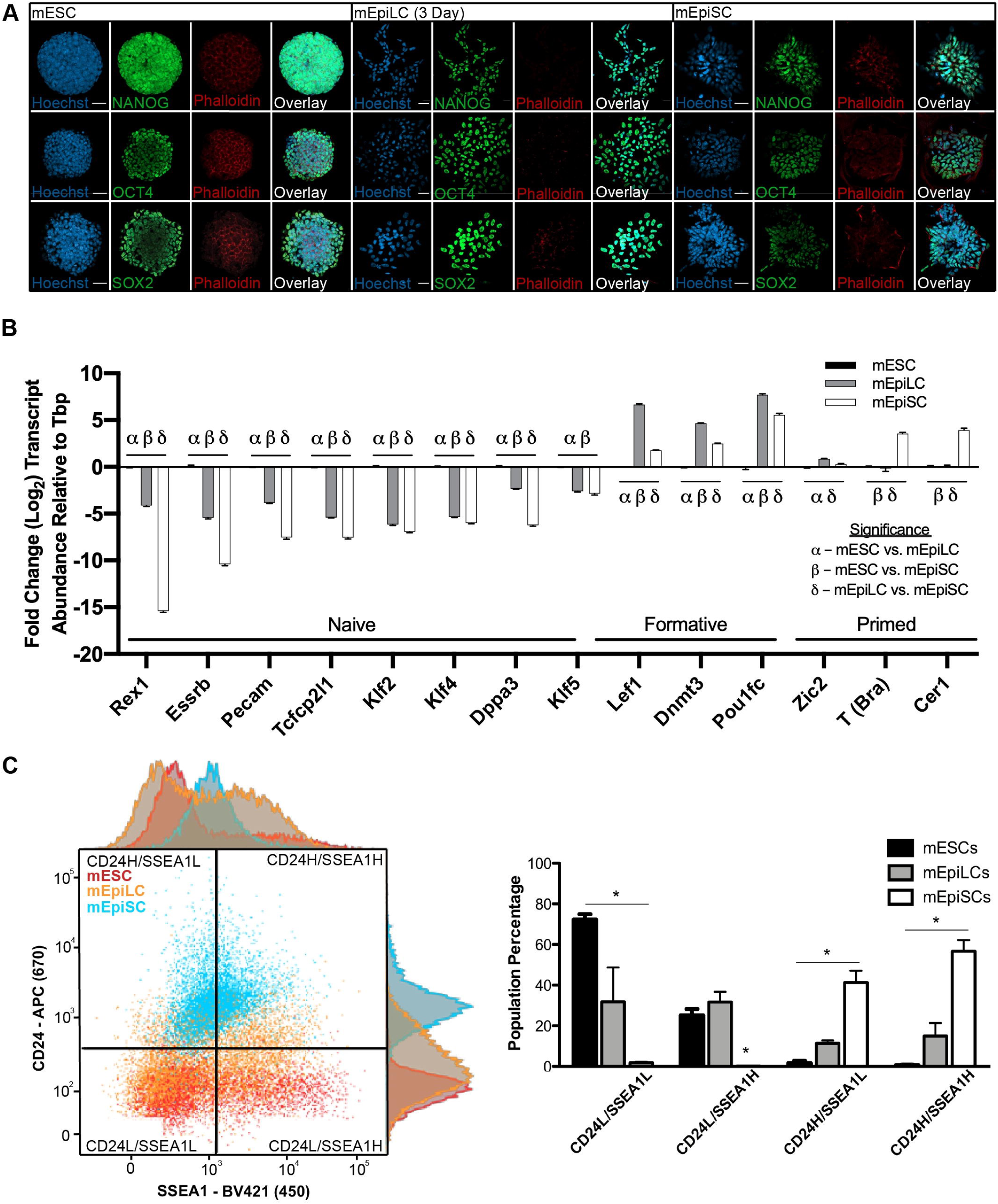
mESC, mEpiLC and mEpiSC populations transcript abundance for pluripotency genes along with CD24 and SSEA1 cell surface marker detection. (A) Immunofluorescence of mESC, mEpiLC and mEpiSC stained with Hoechst (blue), phalloidin (red) and the core pluripotency associated markers (green): NANOG, OCT4, SOX2 assessed by confocal microscopy. Images taken using 40x magnification and scale bars represent 20*μm*. (B) Histogram of transcript abundance of naïve, formative and primed pluripotent associated genes relative to Tbp and normalized to mESCs. Error bars represent standard error of the mean (SEM), n=3, *p<0.05. Statistics for the transcript abundance study represent a two-way ANOVA with Tukey’s multiple comparisons of mean±SEM where ∝=0.05, n=3 biological replicates run in technical triplicate. (C) Histogram depicting flow cytometric analysis of mESCs, mEpiLCs and mEpiSCs comparing presence of the cell surface markers SSEA1 and CD24. Geometric mean is portrayed on both axes of the scatterplot. Error bars represent standard deviation (SD), n=3, *p<0.05. Statistics for the cell surface marker study represent a two-way ANOVA with Tukey’s multiple comparisons of mean±SD where ∝=0.05, n=3 biological replicates run at least in technical triplicate.

Assessment of stage specific transcript abundance of naïve, formative and primed markers verified the pluripotent state of mESCs, mEpiLCs and mEpiSCs, respectively (Fig. 1). The naïve pluripotent associated genes; Rex1, Esrrb, Pecam, Tcfcp2l1, Klf2, Klf4, Dppa3, and Klf5 all underwent a significant (p<0.05) reduction in transcript abundance in mEpiLCs and mEpiSCs relative to mESCs (Fig. 1B). The transcript abundance of formative pluripotent associated genes; Lef1, Dnmt3 and Pou1fc were significantly (p<0.05) increased in mEpiLCs compared to mESCs with Dnmt3 and Pou1fc mRNAs also significantly (p<0.05) elevated in mEpiLCs over that observed in mEpiSCs (Fig. 1B). The transcript abundance of primed pluripotent state associated markers Zic2, T(Brachyury) and Cer1 were significantly (p<0.05) increased in the mEpiSCs relative to the mESCs (Fig. 1B).

Naïve mESCs, formative primed-like mEpiLC and primed mEpiSCs can be distinguished from each other using flow cytometric analysis of cell surface markers (Fig. 1C). Populations of each cell colony were quantified using flow cytometry using detection with the pluripotency associated cell surface markers SSEA1 and CD24. Naïve pluripotency is associated with a positive population of SSEA1 (SSEA1^High^) and negative CD24 (CD24^Low^) cell surface marker expressing cells. We observed a significant (p<0.05) decrease in the number of mEpiSCs with this expression pattern compared to either mESCs or primed-like mEpiLCs. In contrast, primed pluripotency is associated with negative SSEA1 (SSEA1^Low^) and positive CD24 (CD24^High^) cell surface marker expression. mEpiSCs displayed a significantly (p<0.05) greater population of cells with this expression pattern compared to mEpiLCs and mESCs (p<0.05). Unlike the naïve SSEA1^High^ and CD24^Low^ expressing cells, the primed population of SSEA1^Low^ and CD24^High^ expressing mEpiLCs was significantly greater than mESCs (p<0.05). The mEpiLCs cultured 24 hours into the formative pluripotent state (48 hours in FA transitioning media), were still clearly distinguishable from the ‘ground state’, naïve mESCs and the primed mEpiSCs (Fig. 1C).

### PKM1/2 protein abundance and localization fluctuate in naïve mESCs, primed-like mEpiLCs and primed mEpiSCs

We detected a significant (p<0.05) increase in PKM1 and PKM2 protein abundance relative to *β*-ACTIN in formative primed-like mEpiLCs cultured in Fgf-2/Activin A (FA medium) compared to naïve mESCs or primed mEpiSCs (Fig 2A). The ratio of phosphorylated, homodimeric conformation of PKM2 to total PKM2 protein abundance relative to *β*-ACTIN significantly (p<0.05) decreased when naïve mESCs were transitioned to formative primed-like mEpiLCs. However, no significant (p>0.05) difference in the ratio of PKM1 to PKM2 protein abundance relative to *β*-ACTIN was observed in in any pluripotency cell state cultures investigated. PKM1 and PKM2 protein fluorescence were detected in the cytoplasm and nuclei of mESCs, mEpiLCs and mEpiSCs as demonstrated by morphological comparison with Hoechst and rhodamine phalloidin stains representing nuclear and cytoskeletal compartments respectively (Fig 2B). Secondary antibody only controls confirmed the specificity of the PKM1/2 immunofluorescence staining (S3 Fig).

**Fig 2.**
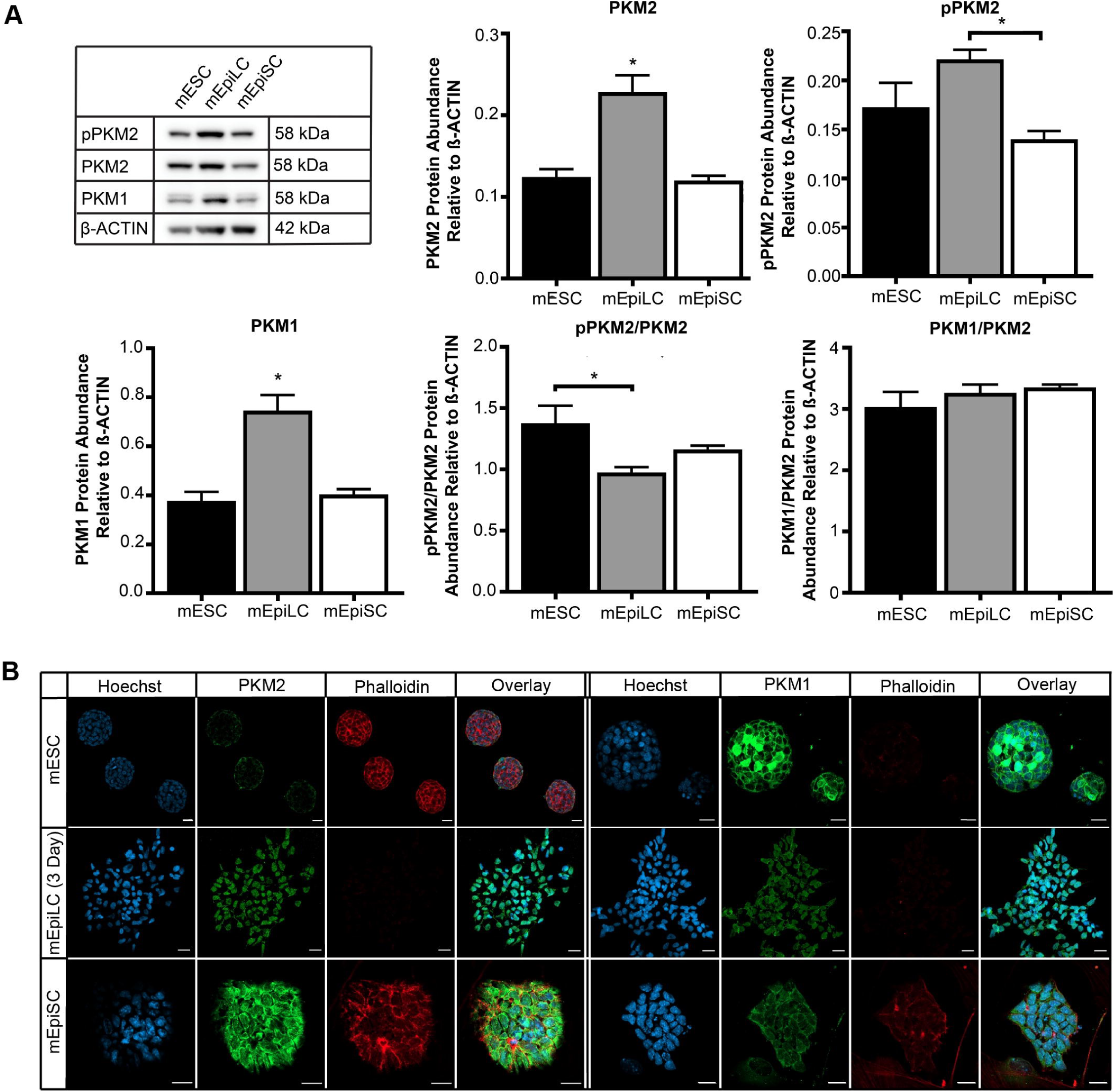
Distinct PKM1 and PKM2 protein profiles in mESCs, mEpiLCs and mEpiSCs. (A) Histogram comparing protein abundance of PKM1, PKM2 and pPKM2 relative to *β*-ACTIN in mESCs, mEpiLCs and mEpiSCs in total protein lysates. Error bars represent SEM, n=3, *p<0.05. Immunofluorescence of mESC, mEpiLC and mEpiSC stained for Hoechst (blue), phalloidin (red) and the metabolic markers: PKM1 and PKM2 (green) assessed by confocal microscopy. Images taken using 40x magnification and scale bars represent 20*μm*. (B) Histogram comparing nuclear, cytoplasmic and total protein lysate abundance of LAMIN A, *α*-TUBULIN, PKM1, PKM2 and pPKM2 relative to total protein lane comparison of Ponceau staining. Error bars represent SEM, n=3, *p<0.05. Statistics represent a one-way ANOVA with Tukey’s multiple comparisons of mean±SEM MOC and PCC scores run in n=3 biological replicates.

### Subnuclear localization of PKM1 and PKM2 with OCT4 within naïve mESCs

To authenticate the subcellular immunofluorescence results (Fig 2B), we performed a colocalization study investigating spatial co-occurrence or overlap and correlation of PKM1 and PKM2 with the nuclear localized marker OCT4 and the cytoplasmic localized marker GAPDH using confocal microscopy. Colocalization of immunofluorescent spatial overlap and correlation was compared using Manders’ overlap coefficient (MOC) and Pearson’s correlation coefficient (PCC), respectively on orthogonal projections with background pixels removed from quantification (63). Using these methods, total mESC colony colocalization of PKM1 and PKM2 with OCT4 and GAPDH showed a high instance of spatial overlap to both marker proteins with a significantly (p<0.05) greater overlap to nuclear OCT4 (Fig. 3A, B). However, PKM1 displayed significantly (p<0.05) higher correlation to OCT4 localization compared to GAPDH (Fig. 4A, B). Using the standards set by Zinchuk *et al.*, PKM1 and PKM2 exhibited a ‘medium’ correlation and a ‘strong’ overlap to both OCT4 and GAPDH localization (PCC range: medium = 0.1-0.48, MOC range: strong = 0.89-0.97) (71). By increasing the signal-to-noise ratio through airyscan processing, the colocalization resolution was improved and the analysis was applied to individual mESCs. Individual cell analysis aligned closely with the colony analysis by indicating a strong correlation for spatial co-occurrence for PKM1 and PKM2 in mESCs (S8A,B Fig). Immunofluorescence controls and colocalization thresholds are shown in S4 Fig.

**Fig 3.**
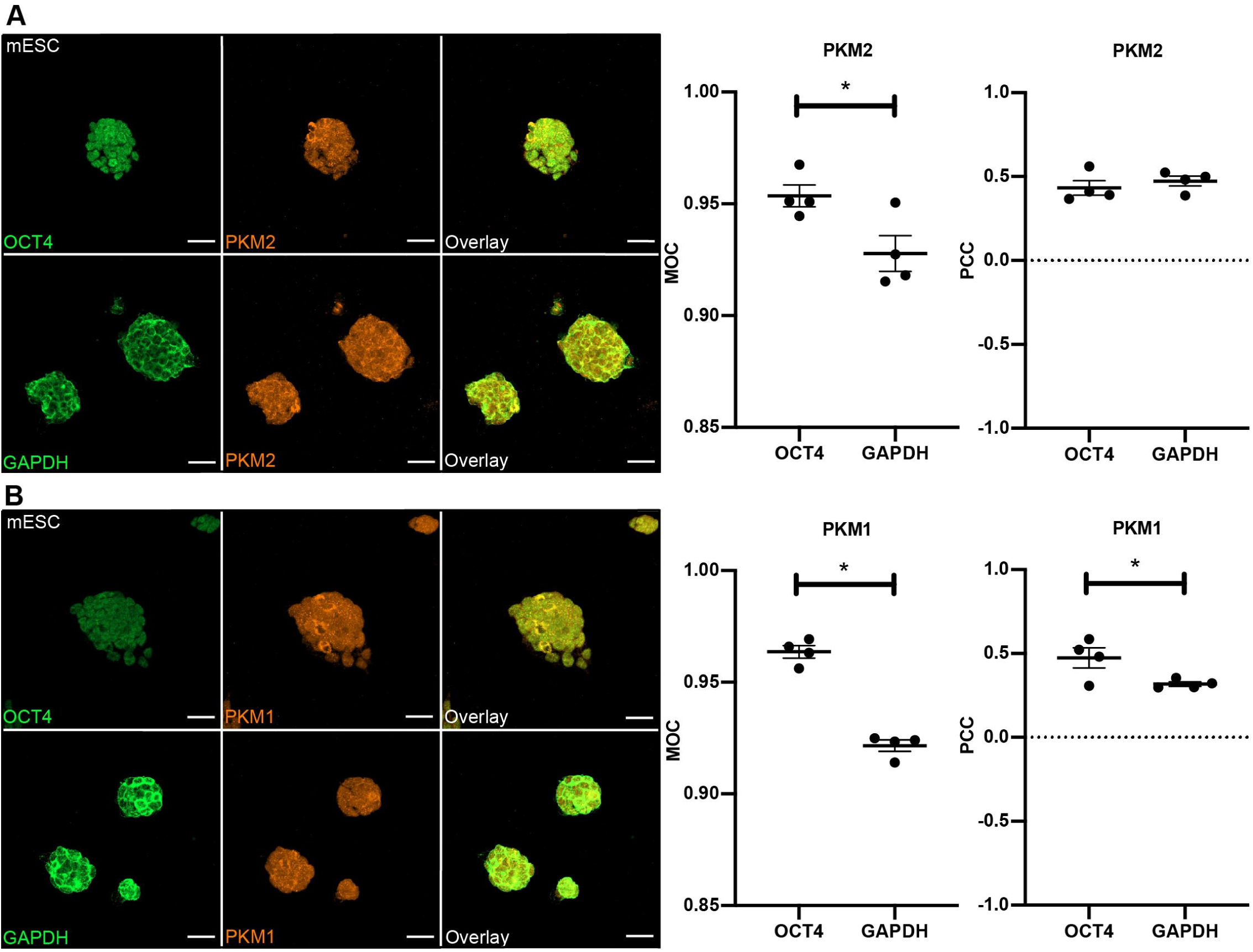
PKM1 and PKM2 are translocated to the nuclei of mESCs and both PKM1 and PKM2 are associated with OCT4 and GAPDH localization. (A) Immunofluorescence of mESCs stained for OCT4 (green), GAPDH (green) and PKM2 (orange) for a confocal, colocalization analysis. Images taken using 40x magnification and scale bars represent 20*μm*. Histogram comparing PKM2 to OCT4 and GAPDH spatial localization by Manders’ Overlap Coefficient (MOC) and Pearson’s Correlation Coefficient (PCC). Error bars represent SEM, n=3, *p<0.05. (B) Immunofluorescence of mESCs stained for OCT4 (green), GAPDH (green) and PKM1 (orange) for a confocal, colocalization analysis. Images taken using 40x magnification and scale bars represent 20*μm*. Histogram comparing PKM1 to OCT4 and GAPDH spatial localization by Manders’ Overlap Coefficient (MOC) and Pearson’s Correlation Coefficient (PCC). Error bars represent SEM, n=3, *p<0.05. Statistics represent a two tailed Mann-Whitney test of mean±SEM MOC and PCC scores run in n=4 biological replicates and at least a technical triplicate.

**Fig 4.**
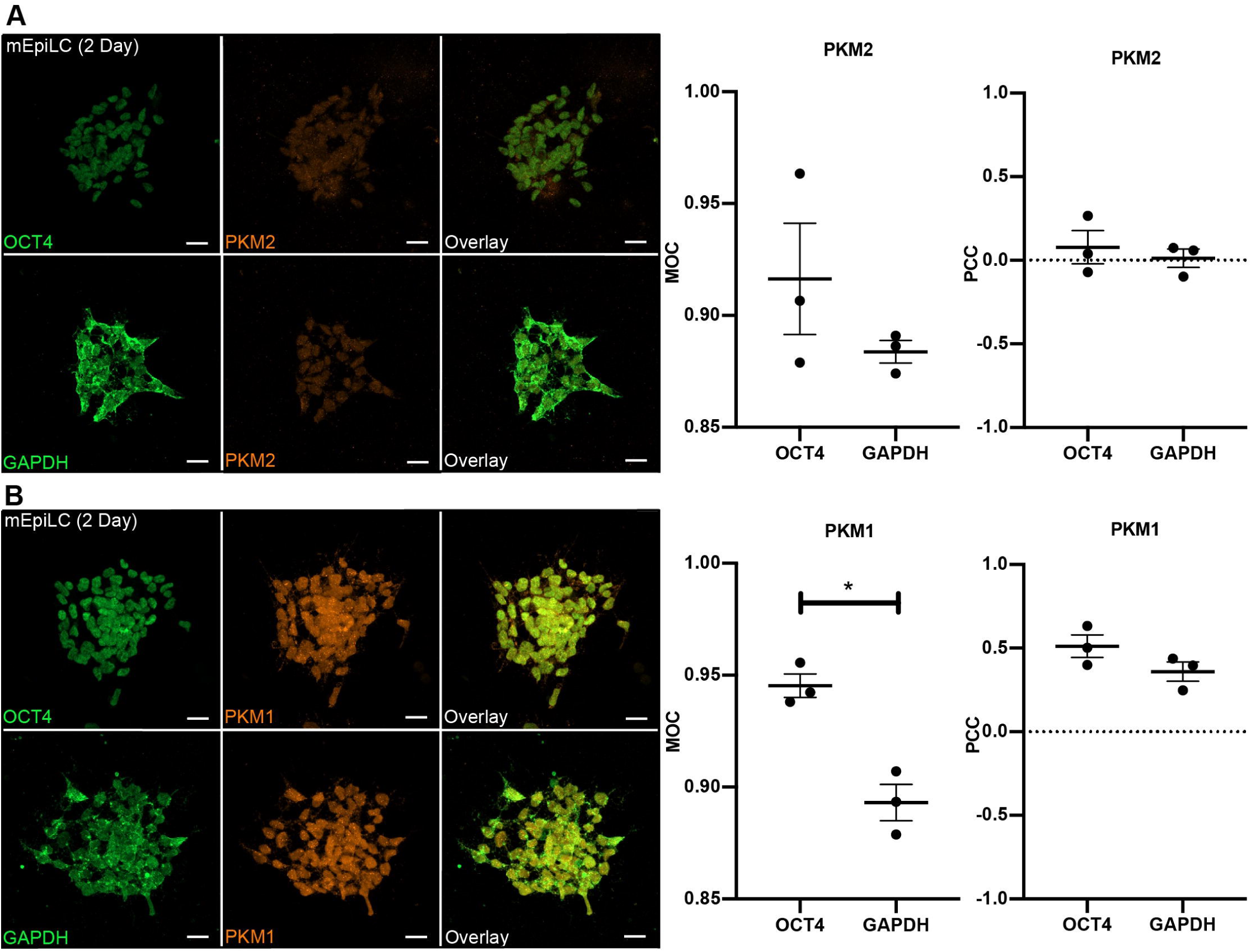
PKM1 and PKM2 are translocated to the nuclei of mEpiLCs and PKM1 is associated with OCT4 and GAPDH localization. (A) Immunofluorescence of mEpiLCs stained for OCT4 (green), GAPDH (green) and PKM2 (orange) for a confocal, colocalization analysis. Images taken using 40x magnification and scale bars represent 20*μm*. Histogram comparing PKM2 to OCT4 and GAPDH spatial localization by Manders’ Overlap Coefficient (MOC) and Pearson’s Correlation Coefficient (PCC). Error bars represent SEM, n=3. (B) Immunofluorescence of mEpiLCs stained for OCT4 (green), GAPDH (green) and PKM1 (orange) for a confocal, colocalization analysis. Images taken using 40x magnification and scale bars represent 20*μm*. Histogram comparing PKM1 to OCT4 and GAPDH spatial localization by Manders’’ Overlap Coefficient (MOC) and Pearson’s Correlation Coefficient (PCC). Error bars represent SEM, n=3, *p<0.05. Statistics represent a two tailed Mann-Whitney test of mean±SEM MOC and PCC scores run in at least n=3 biological replicates and at least technical triplicate.

### Subnuclear localization of PKM1 and PKM2 with Oct4 in mEpiLCs

Immunofluorescent colocalization was quantified in mEpiLCs cultured in transitioning FA medium at 48 hours via confocal microscopy of orthogonal projections and airyscan processing. These cells represent the formative pluripotent state. We applied both total colony and single cell colocalization analysis as described above and observed co-occurrence of PKM1 and PKM2 compared to OCT4 and GAPDH with a significantly (p<0.05) greater PKM1 spatial co-occurrence to OCT4 (Fig. 4A, B). Only PKM1 localization was correlated with both OCT4 and GAPDH localization in these cultures (Fig. 4 B). PKM1 exhibited a ‘strong’ correlation and a ‘strong’ overlap with OCT4 localization, a ‘medium’ correlation and a ‘strong’ overlap to GAPDH localization (PCC range: strong = 0.49-0.84, MOC range: strong = 0.89-0.97) (71).

PKM2 displayed a ‘weak’ correlation to both OCT4 and GAPDH with a ‘strong’ overlap to OCT4 and a ‘medium’ overlap to GAPDH (PCC range: weak = −0.26-0.09, MOC range: medium = 0.71-0.88, strong = 0.89-0.97). Using airyscan processing, individual cells of mEpiLC colonies displayed consistent correlation and spatial overlap compared to the colony as a whole (S9A,B Fig). Immunofluorescence controls and colocalization thresholds are shown in S5 Fig.

### Subnuclear Localization of PKM1 and PKM2 with Oct4 in mEpiSCs

As observed for the naïve mESCs and the formative mEpiLCs, we observed a high degree of PKM1 and PKM2 spatial overlap to both OCT4 and GAPDH in mEpiSCs (Fig. 5A, B). However, unlike the mESCs and mEpiLCs, there were only low levels representing no meaningful correlation of PKM1 or PKM2 with OCT4 or GAPDH in these cultures (Fig. 5A, B). PKM1 and PKM2 immunofluorescence each showed a ‘strong’ overlap to both OCT4 and GAPDH immunolocalizations (MOC range: 0.89-0.97) (71). PKM1 and PKM2 displayed a ‘weak’ correlation to OCT4 and a ‘medium’ correlation to GAPDH (PCC range: weak = −0.26-0.09, medium = 0.1-.48). Using airyscan processing, individual cells of mEpiLC colonies displayed consistent correlation and spatial overlap compared to the colony as a whole (S10A,B Fig). Immunofluorescence controls and colocalization thresholds are shown in S6 Fig.

**Fig 5.**
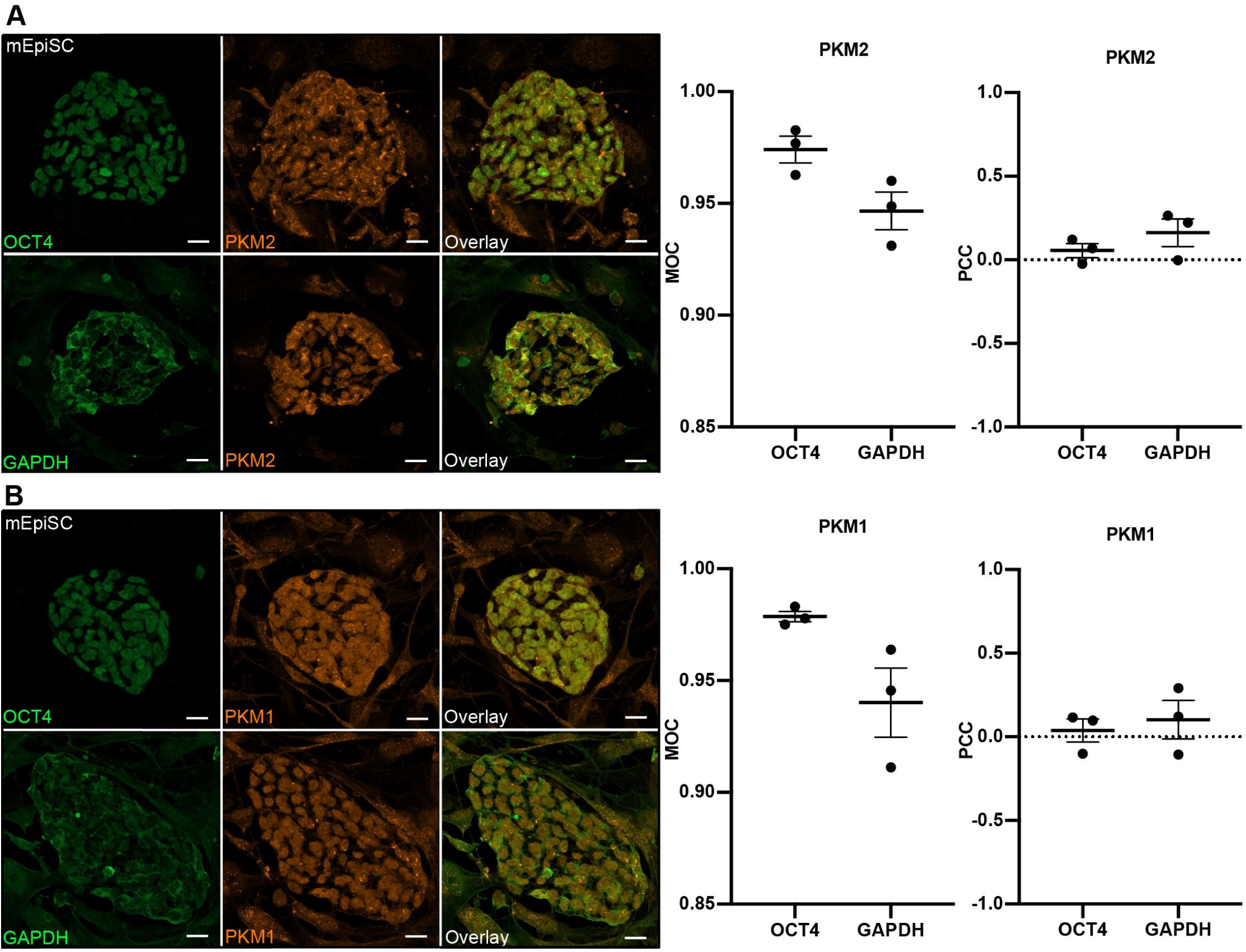
PKM1 and PKM2 are translocated to the nuclei of mEpiSCs and neither isoform is associated with OCT4 or GAPDH localization. (A) Immunofluorescence of mEpiSCs stained for OCT4 (green), GAPDH (green) and PKM2 (orange) for a confocal, colocalization analysis. Images taken using 40x magnification and scale bars represent 20*μm*. Histogram comparing PKM2 to OCT4 and GAPDH spatial localization by Manders’ Overlap Coefficient (MOC) and Pearson’s Correlation Coefficient (PCC). Error bars represent SEM, n=3. (B) Immunofluorescence of mEpiSCs stained for OCT4 (green), GAPDH (green) and PKM1 (orange) for a confocal, colocalization analysis. Images taken using 40x magnification and scale bars represent 20*μm*. Histogram comparing PKM1 to OCT4 and GAPDH spatial localization by Manders’ Overlap Coefficient (MOC) and Pearson’s Correlation Coefficient (PCC). Error bars represent SEM, n=3. Statistics represent a two tailed Mann-Whitney test of mean±SEM MOC and PCC scores run in at least n=3 biological replicates and at least technical triplicate.

### PKM1 and PKM2 are differentially localized to OCT4 and GAPDH between naïve, formative and primed pluripotent states

To obtain a deeper understanding of the cellular co-occurrence of nuclear PKM1 and PKM2 during the transition from mESCs, mEpiLCs and mEpiSCs cultures we contrasted the outcomes between overall co-occurrence (MOC) with Hoechst and OCT4 (positive reference) and Hoechst and GAPDH (negative reference). Relative to the positive reference, there was no significant (p>0.05) changes to MOC of PKM1 or PKM2 localization to OCT4 localization in mESCs, mEpiLCs or mEpiSCs, indicating that PKM1 and PKM2 do indeed occupy nuclear associated regions in these pluripotent cells (Fig. 6B). Relative to the positive reference, there was a no significant (p>0.5) changes to the MOC of PKM1 or PKM2 localization to GAPDH localization in mESCs and mEpiSCs, indicating that PKM1 and PKM2 do indeed occupy cytoplasmic regions in these cells as well (Fig. 6B). However, relative to the positive reference, there was a significant (p<0.05) decrease in MOC of PKM1 and PKM2 localization to GAPDH localization in the mEpiLCs, indicating a decreased cytoplasmic presence in these cells (Fig. 6B).

**Fig 6.**
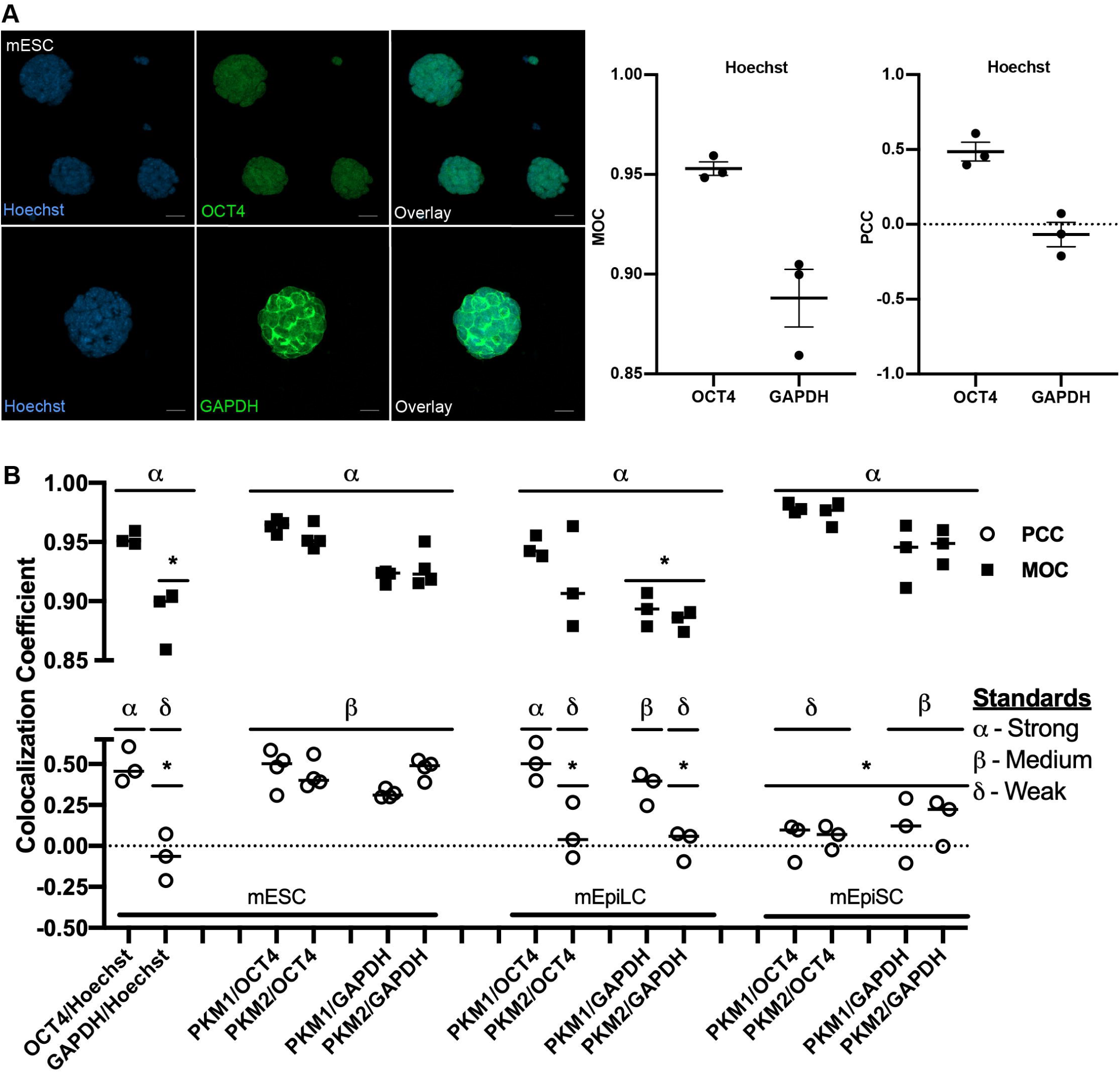
PKM1/2 are moderately associated with OCT4 localization in mESC, PKM1 is strongly associated with OCT4 localization in mEpiLCs and PKM1/2 overlap in nuclear regions of mESCs, mEpiLCs and mEpiSCs. (A) Immunofluorescence of mESCs immuno-stained for OCT4 (green), GAPDH (green) and Hoechst (blue)for a confocal, colocalization analysis. Images taken using 40x magnification and scale bars represent 20*μm*. Histogram comparing Hoechst to OCT4 and GAPDH spatial localization by Manders’ Overlap Coefficient (MOC) and Pearson’s Correlation Coefficient (PCC). Error bars represent SEM, n=3. (B) Total results of colocalization study comparing positive and negative references to mESCs, mEpiLCs and mEpiSC MOC and PCC values. Standard range qualifiers set by Zinchuk *et al*. (2013) compare overlap and correlation differences. Error bars represent SEM, n=3. Statistics of PCC and MOC treatments relative to the positive reference represent a two-way ANOVA with Sidak’s multiple comparisons test of mean±SEM PCC and MOC scores where ∝=0.05, n=3 biological replicates.

To further interrogate the subnuclear association of nuclear PKM1 and PKM2 during transitioning mESCs, mEpiLCs and mEpiSCs cultures, we contrasted the outcomes between overall correlation (PCC) with Hoechst and OCT4 (positive reference) and Hoechst and GAPDH (negative reference). Each mPSC state examined showed differential PKM1/2 subnuclear expression correlation to OCT4 and GAPDH compared to the positive reference. Relative to the positive reference indicating nuclear OCT4 association, there was no significant (p>0.05) difference in PCC of PKM1 or PKM2 localization to OCT4 or GAPDH in mESCs (Fig. 6B). In contrast, mEpiLCs and mEpiSC displayed significantly (p<0.05) less PCC of PKM2 localization to OCT4 relative to the positive reference, however, these values did not reach a meaningful linear correlation level (Fig. 6B). Relative to the positive reference indicating nuclear association, there was no significant (p>0.05) PCC difference in PKM1 and a significant (p<0.05) decrease in correlation of PKM2 localization to OCT4 and GAPDH localization relative to the positive reference in mEpiLCs, suggesting nuclear association of PKM1 and reduced nuclear association of PKM2 with OCT4 (Fig. 6B). Relative to the positive reference indicating nuclear association, there was a significant (p<0.05) decrease in PCC of PKM1 and PKM2 localization to OCT4 and GAPDH localization relative to the positive reference in mEpiSCs (Fig. 6B). However, in the case of mEpiLCs and mEpiSCs, values with PCC = 0 reflect no meaningful linear correlation and we cannot conclusively infer meaningful association of PKM1 or PKM2 localization to these fluorophores of interest.

Using the standard ranges set by Zinchuk *et. al.* to describe these values with qualifying terms, we observed a ‘strong correlation’ and ‘strong overlap’ in the Hoechst/OCT4 positive reference (PCC = 0.49±0.06, MOC = 0.95± 0.00) and a ‘very weak correlation’ and ‘strong overlap’ in the GAPDH/Hoechst negative reference (PCC = −0.07±0.08, MOC =0.89±0.01) (Fig. 6A) (71). These standards promote the superiority of the PCC over the MOC, however, we did see significant differences between our positive and negative references and our sample data indicating a valuable role for the MOC comparison as well.

In summary, PKM1 and PKM2 occupy the same spatial localization as OCT4 nuclear regions and differentially correlate to subnuclear localizations relative to OCT4 and GAPDH localization in mESCs, mEpiLCs and mEpiSCs. We further show that when using the set standard qualifiers of MOC and PCC, the PCC is superior. Additionally, both the PCC and MOC metrics are valuable in comparison to known positive and negative references. Reference stains and colocalization thresholds are available in S7 Fig.

### Nuclear localization of PKM1 and PKM2 in naïve mESCs by cell fractionation

To validate the results of the orthogonal projection immunofluorescence analysis, we conducted a nuclear and cytoplasmic fractionation of naïve mESCs using the REAP protocol (72). Naïve mESCs were selected as they were the only mPSC that exhibited both a nuclear co-occurrence and correlation of PKM1 and PKM2 with OCT4 immunofluorescence from our colocalization study. The REAP protocol was validated by comparing the nuclear and cytoplasmic fractions with the nuclear marker LAMIN A and the cytoplasmic marker ∝-TUBULIN. We observed a significant (p<0.05) increase in ∝-TUBULIN in the cytoplasmic fraction compared to the nuclear fraction validating successful fractionation (Fig. 7A, B, C). The results demonstrated the nuclear localization of PKM1 from naïve mESC protein lysates (Fig. 7 C). This was evident as the ratio of nuclear to cytoplasmic fraction PKM1 trended towards elevated levels of nuclear PKM1 in the mESC, however this did not reach statistical significance (Fig. 7 C) but does agree with the immunofluorescent colocalization outcomes above (Fig. 3B).

**Fig 7.**
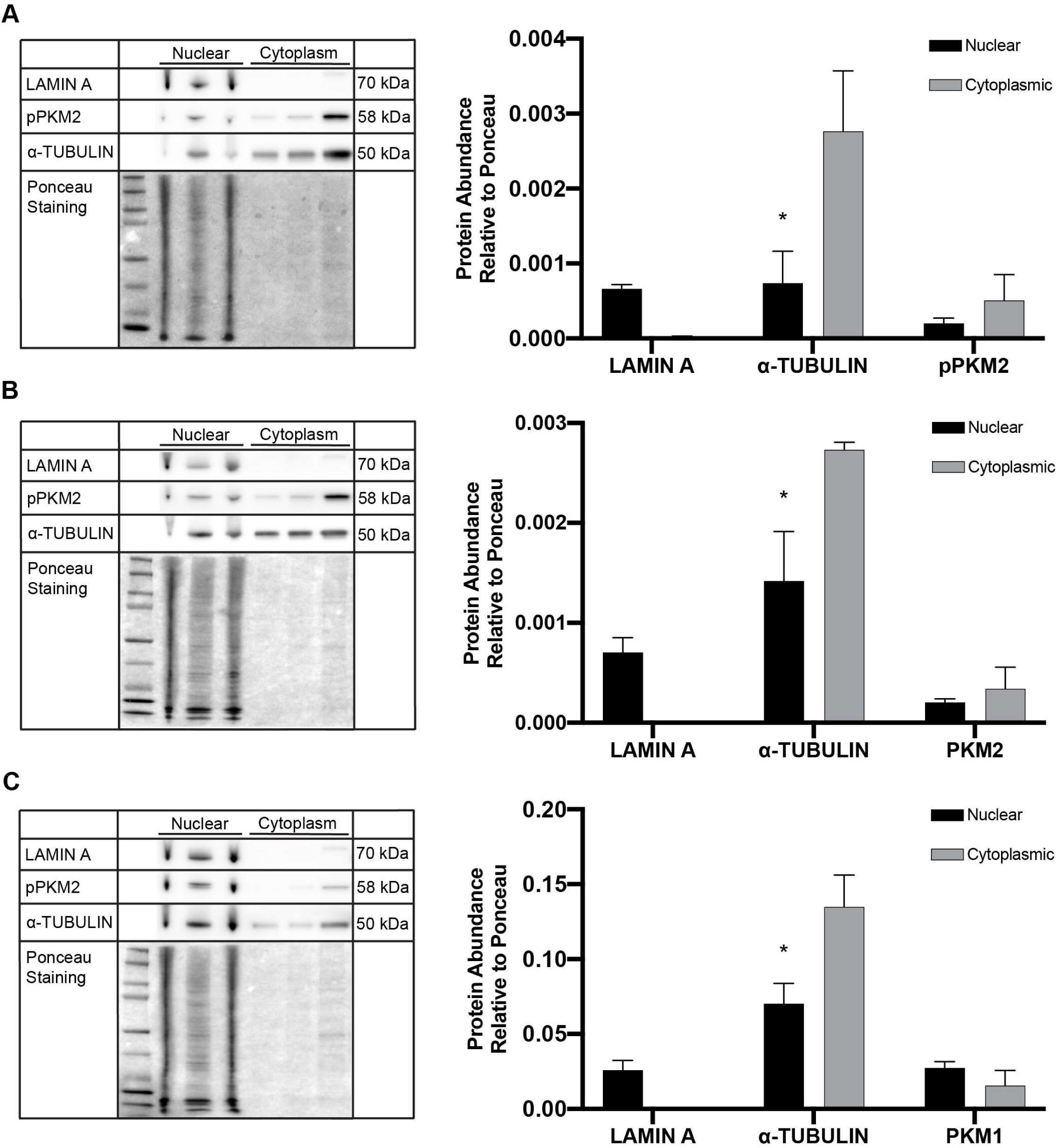
PKM1 is translocated to the nuclei of mESCs. (A) Histogram comparing protein abundance of nuclear and cytoplasmic fractioned lysates of pPKM2, ∝-TUBULIN and LAMIN A relative to total protein Ponceau staining in mESCs. Error bars represent SEM, n=3, *p<0.05. (B) Histogram comparing protein abundance of nuclear and cytoplasmic fractioned lysates of PKM2, ∝-TUBULIN and LAMIN A relative to total protein Ponceau staining in mESCs. Error bars represent SEM, n=3, *p<0.05. (C) Histogram comparing protein abundance of nuclear and cytoplasmic fractioned lysates of PKM1, ∝-TUBULIN and LAMIN A relative to total protein Ponceau staining in mESCs. Error bars represent SEM, n=3, *p<0.05.

## Discussion

Despite traditionally being considered a passive trait of cell-fate determination, mounting evidence now supports metabolism as having a direct role in self-renewal, cell fate and differentiation (35). Our study investigated differences in pyruvate kinase muscle isoforms 1 and 2 (PKM1/2) in naïve, formative and primed pluripotent stem cells and found differential expression and nuclear localization of these metabolic isoforms during pluripotent state transitioning. Densitometry of total protein lysates indicated that over the course of pluripotent progression there is an increase protein abundance of PKM1, PKM2 and phosphorylated PKM2 in the formative state. Despite this increase in protein abundance, the ratio of PKM1 to PKM2, a common ratio used to examine Warburgian cancer cells, was not different between each pluripotent state, indicating that a stable PKM1/2 ratio is likely required for maintaining pluripotency. We observed nuclear immunofluorescence for both PKM1/2 isoforms in naïve mouse embryonic stem cells (mESCs), formative mouse epiblast-like stem cells (mEpiLCs) and primed mouse epiblast stem cells (mEpiSCs). To verify this observation, we devised a confocal colocalization approach to compare differences in nuclear and cytoplasmic localization by contrasting orthogonal projections with well-established reference markers. Using this technique, we determined that in each pluripotent state, PKM1 and PKM2 both reside in nuclear regions and that PKM1 and PKM2 are moderately associated with OCT4 localization patterns in mESCs. PKM1 is strongly associated with OCT4 localization patterns in mEpiLCs and both isoforms have a weak association to OCT4 in mEpiSCs showing a progressive decline in association to the pluripotency gene OCT4 during mouse ES cell pluripotency transitioning. Finally, we demonstrated nuclear translocation of PKM1 in naïve mESCs through nuclear and cytoplasmic fractionation using the recently described REAP protocol of cell fractionation (72). We also applied a new flow cytometry method to distinguish the formative state pluripotent state using the cell surface markers CD24 and SSEA1 and devised an effective colocalization strategy for nuclear and cytoplasmic immunolocalization quantification, addressing the misconceptions regarding Manders’ overlap coefficient (MOC) and Pearson’s correlation coefficient (PCC) that exist in the recent literature (69,70).

Our initial work highlights the usefulness of quantifying the expression of the cell surface markers CD24 and SSEA1 via flow cytometry to distinguish different pluripotent populations. This technique allowed us to evaluate the incidence and degree of mEpiLCs, exhibiting the formative pluripotent state, that had exited naïve mESCs and transitioned toward primed pluripotency. This method of using CD24 and SSEA1 has been used to characterize naïve and primed cells (73), but this is the first time that transitioning cells have been compared to either end of the pluripotent continuum. To validate the transitioning from naïve to formative pluripotency, quantitative RT-PCR revealed the high abundance of formative associated transcripts; Dnmt3, Lef1 and Pou1fc relative to the naïve and primed states (20). This flow cytometry technique should be useful in FACS sorting and evaluating mEpiLCs along with cells of alternate and region-selective pluripotent states (2,74). As these cells are the only PSCs capable of efficiently differentiating into primordial germ cell-like cells (PGCLCs), this protocol could be useful for the study of germ cell differentiation (2,14).

The measurement of colocalization is a complicated and hotly debated area of biology (69,70). The term colocalization is largely used to measure two main components with different applications, namely correlation or co-occurrence of two fluorophores to each other based on pixel distribution (66). Co-occurrence in immunofluorescence is the presentation of fluorescent pixels existing in the same spatial distribution, and it is an indicator of overlap between markers (65). Correlation is a measurement of the relationship between the pixel intensities and may indicate a biochemical interaction (65). Both the Manders’ overlap coefficient (MOC) and Pearson’s correlation coefficient (PCC) are valid measures of colocalization but they inform different biological questions (70). Immunofluorescence is commonly thought of as a qualitative technique and the literature into colocalization often uses descriptors such as moderate or strong association within PCC ranges. Zinchuk *et al.* (71) developed a method of colocalization range descriptors to bring greater consistency to the field and offer more validity to the quantitative nature of colocalization. We implemented this approach to assign a quantitative designate to the colocalization of PKM1 and PKM2 within the mPSCs of this study. Our study supports claims that the MOC is a valuable metric of colocalization. By comparing MOC and PCC values to a positive and negative biological reference, we were able to set a stronger baseline than using only improved descriptors. We used well recognized nuclear (OCT4) and cytoplasmic (GAPDH) proteins as control markers to compare to another known nuclear stain, Hoechst, which set a positive and negative reference to nuclear colocalization that allowed us to directly compare MOC and PCC values to. Comparing our known positive and negative references to the qualifying range standards set by Zinchuk *et al*. our data supports that comparing colocalization by correlation is superior to spatial overlap in our system (71). However, while MOC still provided valuable knowledge, the PCC data showed an improved distinction between internal reference controls. Our findings demonstrate that it is critical to run positive and negative references relative to dual fluorophore colocalization and that in the case of mouse embryonic stem cells, the spatial overlap data may not be sufficient to reach quality colocalization assessment compared to correlation data when considering the qualifying standards set by Zinchuk *et al*. We observed that the MOC metric in mPSCs did not delineate nuclear and cytoplasmic distinctions by colocalization and that the PCC metric was a highly effective and viable tool for such distinction and analysis. To increase the power of our colocalization study, we did not simply analyze single images but employed orthogonal projections of stacks examining the data of individual slices to characterize the localization patterns of a true three-dimensional structure. We also accounted for the inherent flaws of the MOC calculation by examining only the individual colonies and individual cells in the orthogonal and airyscanned images respectively to prevent autofluorescence or background pixel offset to influence the algorithm.

Naïve mESCs, in the metabolically bivalent state, proved to be a unique and attractive cell type for colocalization analysis. By examining the correlation of PKM1 and PKM2 immunolocalizations to OCT4 and GAPDH immunolocalizations, we were able to assess not only if the PKM isoforms were occupying similar spaces, but if the trends in subnuclear pixel intensity were related as well. Not only did both isoforms occupy the same spatial regions in comparison to the controls, but both PKM1 and PKM2 were clearly associated with the localization patterns of both OCT4 and GAPDH. Together, these results promote the concept that PKM1 and PKM2 both translocate to the nuclei of mESCs. A recent study using mass spectroscopy of human lung carcinoma cells determined that PKM1 and PKM2 interact with each other (60) suggesting a possible PKM1/2 interaction in the nuclei of mouse ESCs. To validate our colocalization study, we examined the PKM1/2 protein abundance in nuclear and cytoplasmic fractions of mESCs. Due to the inherent difficulty of nuclear and cytoplasmic extraction and the exceptionally high nuclear-to-cytoplasmic ratio of mESCs, we eventually found a successful method of extraction, the REAP or Rapid, Efficient and Practical method of extraction (72,75). Using this technique, we were able to determine that PKM1 and PKM2 do exhibit nuclear lysate protein abundance and that PKM1 is enriched in the nuclear fraction compared to the cytoplasmic fraction of mESCs, supporting our novel observation that PKM1 is translocated to the nuclei of naïve mESCs.

In our initial PKM protein abundance characterization of total cell lysate we found that there was an increase of PKM1 and PKM2 levels in mEpiLCs. Despite this increase in protein abundance, the ratio of PKM1 to PKM2 protein abundance did not change between any of the pluripotent cell types examined. As PKM2 switches to increased PKM1 expression during differentiation and development, with the reverse occurring during tumor formation, the role of the PKM1 to PKM2 ratio has become a focus of interest (76). It may be more pertinent to examine the nuclear to cytoplasmic ratio of PKM1/2 including the dimer to tetramer conformations of PKM2 in various pluripotent states. Surprisingly, the formative state mEpiLCs were significantly different in PKM1 or PKM2 colocalization spatial overlap to OCT4 compared with the positive reference. This demonstrates very low amounts of either isoform occupying the traditional cytoplasmic region occupied by GAPDH for both isoforms. When examining mEpiLCs for correlation of PKM1 and PKM2 colocalization to OCT4 and GAPDH, we determined that PKM1 was associated with both OCT4 and GAPDH compared to the controls. Coupling this finding with the results of the colocalization overlap findings, the formative state mEpiLCs were unique in primarily localizing PKM1 in the nucleus, suggesting that PKM1 may be key in the transition of bivalent metabolism to preferential aerobic glycolysis. Previous studies have shown that the transcription factor promyelocytic leukemia protein (PML), a known PKM2 mediator that maintains the homotetrameric conformation and suppresses the Warburg effect, interacts with OCT4 and NANOG and is necessary for maintaining naïve pluripotency (77–79). Knocking down or deleting PML resulted in flat, slower growing mESC colonies with reduced OCT4, SOX2, cMYC and NR0B1 and diminished naïve-associated BMP, LIF/STAT3 and PI3K signaling whereas Activin A and FGF signalling increased (77). Overexpression of PML resists mESC transitioning towards primed pluripotency and is required for efficient iPSC generation (77). Future studies should examine the influence of PML in the generation of formative state mEpiLCs. As mEpiLCs are the only cells currently described that can efficiently give rise to primordial germ-like cells, PML and PKM1/2 may be important targets for controlling cell fate to efficiently produce mEpiLCs (13).

Finally, our colocalization study of mEpiSCs was also quite revealing. We determined that of all the mPSCs we studied, the primed mEpiSCs had the greatest spatial overlap as assessed by Manders’ overlap coefficient (MOC) of PKM1 and PKM2 colocalization to OCT4 and GAPDH yet significantly lower PKM1 and PKM2 correlation (PCC) to OCT4 and GAPDH. This was somewhat surprising as other Warburgian cells such as glioma stem cells display an interaction between PKM2 and OCT4 (52). The reduced association as assessed by the Pearson correlation coefficient (PCC) of PKM2 and OCT4 may reflect differential chromatin targets in the primed pluripotent state and may be associated with lineage priming and reduced differentiation potential (52). Interestingly, there is also a decrease in PKM1 correlation to OCT4 as assessed by PCC, but only in the primed mEpiSCs. Using our refined colocalization analysis we show that PKM1 and PKM2 co-occur (MOC) in the nuclei of mPSCs across the pluripotent continuum and that PKM1 and PKM2 are differentially correlated (PCC) with OCT4 and GAPDH in each examined pluripotent state. Our findings suggest that ChIP-sequencing of PKM1 and PKM2 targets should be examined in mPSC varieties encompassing the pluripotent continuum. Further, the correlation of PKM2 colocalization to OCT4 decreases from naïvety through the formative state and into primed pluripotency. As such, we conclude that nuclear PKM1 and PKM2 are implicated as contributors to the maintenance and progression of early stem cell pluripotency.

Recent literature has reported instances of nuclear and mitochondrial translocation of PKM2 (80,81). The nuclear translocation of PKM2 is implicated in the regulation of the master glycolysis regulator HIF-1α (56). Jumonji C Domain-containing dioxygenase 5 (JMJD5)-PKM2 interaction hinders PKM2 tetramer formation, blocks pyruvate kinase activity and promotes translocation of PKM2 into the nucleus to regulate HIF-1α-mediated gene transcription (56). JMJD5 regulates the cell cycle and maintains pluripotency in human embryonic stem cells (82), however its role in the nuclear translocation of PKM2 and regulating metabolism in pluripotent stem cells has not been explored. Overexpression of PKM2 maintains the undifferentiated state by fine tuning redox control in naïve mESCs grown as embryoid bodies (83). Future studies treating naïve stem cells with pharmacological agents such as shikonin or DASA-58, which promote the tetrameric conformation of PKM2, may resist formative state transitioning by maintaining the naïve state (61,84). Adjusting PKM2 levels has been completed in mESCs and a complete knockout should be feasible as PKM2-null mice are viable though they experience some metabolic distress and have a reliance on PKM1 (85). However, these mice show induction of late onset formation of spontaneous hepatocellular carcinomas (85). PKM2 is certainly a potential target for cancer treatments and likely a key player in cellular reprogramming and differentiation (36,85). Despite several non-canonical roles being characterized, it is likely that other roles exist and have yet to be discovered (57).

While PKM2 has been extensively studied in cancers and stem cells (76,83,85–89), the PKM1 isoform has not been investigated to the same extent. There is a growing body of evidence to suggest that PKM1 may play an important role in early differentiation and within specific cancer subtypes. PKM1 is essential for the proliferation and tumor-promoting capabilities of small cell lung cancer (SCLCs) and other net endocrine tumors (76). Oxygen consumption in PKM1 overexpressed cancer cells does not change although there are more mitochondria with a greater rate of mitochondria dysfunction, while there are more reactive oxygen species generated in the PKM2 overexpressed cells compared to the PKM1 overexpressed cells (76). These characteristics of PKM1 overexpressed cells are accompanied with increased autophagic flux and increased tumor growth with increased autophagy and mitophagy (76). PKM1 could play a non-canonical role in promoting autophagic and mitophagic roles during pluripotent stem cell state transitioning. When either PKM1 or PKM2 was overexpressed in mESCs, it was found that the pluripotency markers Nanog, Eras and Rex1 were upregulated and an embryoid body formation assay showed that overexpression did not influence differentiation (25). Taken together, these results indicate that PKM1 contributes to proliferation, stemness and pluripotency. Based on our protein abundance analysis PKM2 or both isoforms may promote the generation of mEpiLCs and the formative pluripotent state (36). Our results suggest that the ratio of PKM1 to PKM2 may be necessary to maintain mouse pluripotency. We also report a unique localization of PKM1 that suggests a novel, non-canonical role just as nuclear, dimeric pPKM2 has been implicated in several non-metabolic roles associated with stemness and cell growth (48). Recently, the role of PKM1 in highly proliferative cells has been highlighted (76). These results along with our current data questions PKM2’s role as the traditional prototypic isozyme of development as it is now clear that PKM1 is expressed and likely has non-canonical roles (76). Nuclear PKM1 has recently been reported in other highly proliferative cell types such as human liver cancer cells (HepG2 and SMMC-7721) (55). Following treatment with drug Oroxylin A (OA), PKM1 is translocated to the nucleus with hepatocyte nuclear factor 4 α (HNF4α) and increases the PKM1 to PKM2 ratio resulting in hepatoma differentiation (55). PKM1 overexpressed in embryoid bodies generated from mESCs resulted in increased endoderm transcript abundance of FOXA2, AFP and HINF1B, implicating PKM1 in endoderm differentiation (83). Given our colocalization findings, the nuclear localization of PKM1is certainly implicated in formative state generation and the addition of a drug such as OA may modulate the occurrence of this transient pluripotent state.

In summary, we have demonstrated differential nuclear and subnuclear localization of both PKM1 and PKM2 in mouse pluripotent stem cells and suggest a novel regulatory role for nuclear PKM1. We have established differential nuclear, subnuclear and cytoplasmic association of PKM1 and PKM2 in mESC cells as they transition from naïve pluripotency, through formative state (primed-like mEpiLCs) towards primed mEpiSCs. We suggest that protein colocalization studies applied to PSCs should give greater weight to their correlation data and not their spatial overlap findings especially if the standards set by Zinchuk *et al.* are implemented (71). The presence of nuclear PKM1/2 and the dynamic redistribution of PKM1 and PKM2 during pluripotency continuum suggests potential non-canonical roles for both isoforms in maintaining and directing varying pluripotent states.

## Materials and Methods

### Antibody specificity

Rabbit polyclonal antibodies specific for PKM1 and PKM2 (Proteintech 15821-1-AP, 15822-1-AP) were used to distinguish between PKM1 and PKM2 protein localization and abundance in this study. These PKM1 and PKM2 antibodies recognize the corresponding immunogens of LVRASSHSTDLMEAMAMGSV and LRRLAPITSDPTEATAVGAV, respectively, and have been knockdown-verified confirming their isoform specificity (38,90–93).

### Feeder cell derivation and culture conditions

Mouse embryonic fibroblasts (MEFs) (CF1 cell line, ThermoFisher) derived from E12.5 mouse embryos were plated and expanded on 0.1% porcine gelatin (Sigma G2500) coated dishes and irradiated (8000 rads). Irradiated MEFs were cultured in media containing the following; DMEM (ThermoFisher11960044), 8.9% Qualified FBS (ThermoFisher 12483020, lot# 1936657), 1.1% MEM NEAA (100x) (ThermoFisher 11140050), 1.1% GlutaMAX (ThermoFisher 35050061). Irradiated MEFs were plated on 0.1% gelatin dishes and cultured for a minimum of 1 hour prior to mEpiSC plating for immunofluorescence and 24 hours for all other molecular analyses.

### Stem cell culture conditions

Feeder-free, naïve, mouse embryonic stem cells (mESCs, R1 strain – 129X1 x 129S1; provided courtesy of Dr. Janet Rossant, Hospital for Sick Children, Toronto, Canada), feeder-free, primed-like mouse epiblast-like cells (mEpiLCs, chemically converted R1 mESCs) and primed mouse epiblast stem cells (mEpiSCs, strain – 129S2; also provided by Dr. Janet Rossant) were cultured in the following base media; a 1:1 mixture of KnockOut DMEM/F12 (ThermoFisher 12660012) and Neurobasal Media (ThermoFisher 21103049) with 0.1% 2-Mercaptoethanol (Gibco 21985-029), 0.25% GlutaMAX (ThermoFisher 35050061), 1.0% N2 Supplement (100x) (ThermoFisher 17502048), 2.0% B27 Supplement (50x) (ThermoFisher 17504044). Base media for the culture of mESCs were supplemented with 1000 units/mL ESGRO Recombinant mouse LIF protein (EMD Millipore ESG1107), and 2i small molecule inhibitors: 1μM PD0325901 (Reagents Direct 39-C68) and 3μM CHIR99021 (Reagents Direct 27-H76). Base media for the culture of mEpiLCs and mEpiSCs were supplemented with 20ng/mL Activin A from mouse (Sigma-Aldrich SRP6057) and 12ng/mL Fgf-2 from mouse (Sigma-Aldrich SRP3038). mESCs were passaged using Accutase (STEMCELL Technologies 07920) and centrifuged at 244 x g for 5 minutes. Primed mouse epiblast stem cells were cultured in the base medium and supplements as mEpiLCs were along with a substratum of irradiated mouse embryonic fibroblasts (MEFs).

One hour prior to passaging, growth medium was replaced. Passaging was completed using Gentle Cell Dissociation Buffer (Gibco 13151-014) for 5 minutes at room temperature. Lifted cells were then centrifuged at 244 x g for 3 minutes and plated at a density of 1:12 onto fibronectin coated dishes with MEFs. RNA and protein abundance studies were completed by excluding MEFS for feeder-free conditions and passaging mEpiSCs with StemPro™ Accutase™ (Thermo Fisher A1110501) to ensure only MEF free lysates were used. In the case of the flow cytometry experiments, a size-exclusion gating strategy and the presence of pluripotency markers was used to delineate between MEF and mEpiSC populations. Additionally, during the preliminary work for this study it was clear that the MEF feeder cells supporting the mEpiSCs, express both PKM isoforms in abundance. We weaned our mEpiSCs off irradiated MEFs by gentle enzymatic passaging onto fibronectin over two passages, this resulted in a clean and healthy population of feeder-free mEpiSCs ready for protein abundance studies.

### Real-Time Quantitative qRT-PCR

RNA isolation was completed using a RNeasy RNA isolation kit (Qiagen 74104) and Trizol (Ambion 15596018) hybrid protocol followed by DNAse treatment (Invitrogen AM1906). cDNA synthesis was completed in accordance to iScript (BioRad 170-8891) protocols. Total RNA was extracted from adherent cells using TRIzol Reagent (Invitrogen) and a RNeasy mini kit (Qiagen). DNAses were then removed using DNAse Free Kit (AM1906). cDNA synthesis using iScript. Primers were tested in temperature gradients before cDNA dilution series to determine primer efficiencies. Relative transcript abundance was compared using mean±SEM with a two-way ANOVA with Tukey’s multiple comparison test with three biological replicates. Relative transcript abundance was calculated using the Pfaffl method of quantification, normalized to mESCs and relative to TATA-binding protein (Tbp) transcript abundance. Forward and reverse primer designs and annealing temperatures are available in Table 1.

**Table 1.**
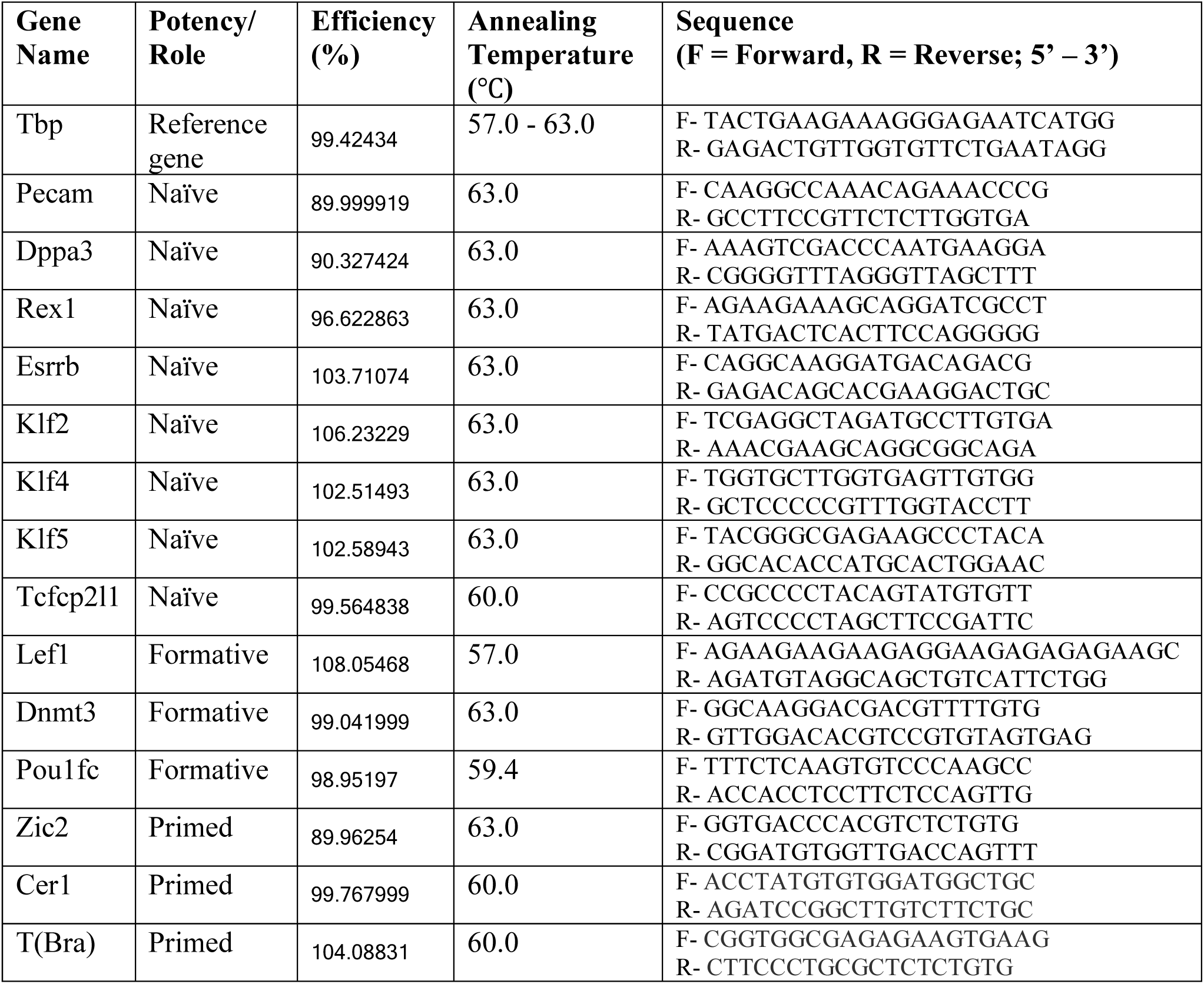
PCR Primers.

### Western blotting

Mouse ESCs and mEpiLCs were washed with cold DPBS (calcium chloride/magnesium chloride/) (PBS(+/+)) (Gibco 14040-133) and all cell types were lysed with Pierce™ RIPA Buffer (ThermoScientific 89900) supplemented with 1X Phosphatase Inhibitor Cocktail Set 2 (Calbiochem 5246251) and 1X Protease Inhibitor Cocktail Set 1 (Calbiochem 539131). mEpiSCs passaged off MEF-coated plates onto fibronectin (Roche 11051407001) coated plates for a single passage using StemPro™ Accutase™ to avoid MEF contamination. mEpiSCs were centrifuged at 244 x g and lysed. Lysates were sonicated for five, 30 joule pulses over 30 seconds and were rotated at 4°C for 30 minutes followed by centrifugation of 12000 rcf at 4°C for 20 minutes with the supernatant removed into a fresh, chilled tube. Protein quantification was completed using a Pierce™ BCA Protein Assay kit (Thermo Fisher Scientific 23225). Protein loading mixes were prepared at 10-25ug samples in MilliQ H_2_O, LDS and Reducing Agent at 70°C for 10 minutes. Loading mixes were loaded in NuPAGE™ 4-12% Bis-Tris Gels (Invitrogen NuPAGE NP0336), the electroporation solution consisting of 1x MOPS (BOLT Invitrogen B000102) and 500uL of sample reducing agent containing dithiothreitol (Thermofisher NP0009) added. Electrophoresis was completed at 200V for 50 minutes. Proteins were transferred to a PVDF membrane at 100V for 2 hours in ice-cold conditions. The protein transferred PVDF membrane (EMD Immobilon IPVH00010) was blocked in 5% bovine serum albumin (BSA) (ALB001) for phosphorylated antibodies or 5% skimmed milk (Carnation) in 1x TBST for 1 hour at room temperature with end-to-end agitation. Primary antibodies were introduced to the membrane overnight at 4°C with rotation. Membranes were washed 3 times for 10 minutes in TBST and HRP conjugated secondary antibodies were introduced for 1 hour at room temperature with rotation. Membranes were then washed three times for 10 minutes each and imaged with Luminata Classico Western HRP Substrate (EMD WBLUC0500) or Immobilon Forte Western HRP Substrate (EMD WBLUF0500) on a ChemiDoc. Membranes were stripped using antibody stripping buffer (FroggaBio ST010) until previous antibody binding was no longer evident. Bands of interest were compared to *β*-ACTIN and/or Ponceaus S for total lane protein densitometry. Western blotting densitometry results were compared using mean ± SEM, One-way ANOVA, Tukey’s multiple comparison test with three biological replicates. Primary and secondary antibodies and their concentrations are listed in Table 2.

**Table 2.** Western blot antibody/marker list.

| **Primary Antibody/Marker** | **Concentration** | **Secondary Antibody** | **Concentration** |
| --- | --- | --- | --- |
| β-ACTIN<br>A3854 | 1:50000 | N/A: HRP-linked | N/A: HRP-linked |
| PKM1<br>15821-1-AP | 1:5000,<br>5% TBST | Anti-rabbit IgG,<br>HRP-linked<br>Antibody #7074<br>Cell Signaling<br>Technology | 1:2000-10,000,<br>5% Milk in<br>TBST |
| pPKM2<br>(Tyr105)<br>3827S | 1:5000,<br>5% BSA in<br>TBST | Anti-rabbit IgG,<br>HRP-linked<br>Antibody #7074<br>Cell Signaling<br>Technology | 1:2000-10,000,<br>5% BSA in<br>TBST |
| PKM2<br>15822-1-AP | 1:5000,<br>5% Skim milk<br>in TBST | Anti-rabbit IgG,<br>HRP-linked<br>Antibody #7074<br>Cell Signaling<br>Technology | 1:2000<br>5% Milk in<br>TBST |
| OCT4<br>sc-5279 | 1:5000<br>5% Skim milk<br>in TBST | Donkey anti-<br>mouse IgG-HRP<br>sc-2314<br>Santa Cruz | 1:10,000,<br>5% Skim milk<br>in TBST |
| GAPDH<br>sc-32233 | 1:10000<br>5% Skim milk<br>in TBST | Donkey anti-<br>mouse IgG-HRP<br>sc-2314<br>Santa Cruz | 1:10,000,<br>5% Skim milk<br>in TBST |
| LAMIN A<br>Ab26300 | 1:10000<br>5% Skim milk<br>in TBST | Anti-rabbit IgG,<br>HRP-linked<br>Antibody #7074<br>Cell Signaling<br>Technology | 1:2000<br>5% Skim milk<br>in TBST |
| α-TUBULIN | 1ug/mL<br>5% Skim milk<br>in TBST | Anti-rabbit IgG,<br>HRP-linked<br>Antibody #7074<br>Cell Signaling<br>Technology | 1:2000<br>5% Skim milk<br>in TBST |
| Ponceau Stain | 0.1% (w/v), 5%<br>acetic acid | N/A | N/A |

### Flow cytometry

Each cell type; mESCs, mEpiLC (3 days of conversion), and mEpiSCs were lifted with TrypLE (Gibco 12605-028) for 5 minutes at 37°C and inactivated in MEF culture medium. This single cell suspension was then spun down at 244 x g for 5 minutes. Dead cell gating was accomplished by taking a proportion of approximately 5% of each cell type and pooling them into a single treatment and killed at 60°C for 10 minutes. Live and dead treatments were resuspended in 100uL of PBS and stained with 1:1000 zombie aqua dye (excluding appropriate full minus one controls) and incubated in the dark at room temperature for 30 minutes. Cells were washed with 2mL of flow cytometry staining buffer (FCSB) containing: 97% PBS (-/-), 3% FBS (qualified, ESC grade), and centrifuged at 244 x g for 5 minutes prior to fixation with 10% paraformaldyde (PFA) in PBS for 10 minutes. Fixed cells were washed with PBS, centrifuged before being split into individual treatments for staining. Allophycocyanin (APC) and phycoerythrin (PE) antibodies were added and vortexed for 1 hour in the dark at room temperature prior to washing, centrifugation and resuspension in 200 uL of PBS before being ejected through a 40 μM cell strainer (Falcon 352340) and final wash with 300 μL of PBS. Compensation beads were stained for each marker (excluding zombie stain) for 30 minutes at room temperature in the dark and washed with 2 mL of FCSB. The beads were centrifuged at 244 x g for 5 minutes and reconstituted in FCSB. Flow cytometry was completed on a FACSCanto™ flow cytometer. Flow cytometry population events were compared using mean±SD with a one-way ANOVA with Tukey’s multiple comparison test for unpaired, two tailed t-test respectively with three biological replicates. Antibodies and their concentrations are listed in Table 3.

**Table 3.** Flow cytometry antibody/marker list.

| Primary Antibody/Marker | Concentration | Secondary Antibody | Concentration |
| --- | --- | --- | --- |
| CD24<br>138505 | 0.4ug/million cells | N/A:<br>Conjugated | N/A:<br>Conjugated |
| SSEA1<br>125614 | 5ul/million cells | N/A:<br>Conjugated | N/A:<br>Conjugated |

### Immunofluorescence and confocal microscopy

Cells were plated onto 1.25mm thick coverslips coated with gelatin. When ready, cells were fixed with 2% paraformaldehyde (PFA) (EMS 15714) in PBS(+/+) for 10 minutes and washed for 5 minutes with chilled PBS(+/+). Following fixation, cells were permeabilized with 0.1% Triton X-100 (TX1568-1) in PBS(+/+) for 10 minutes and washed for 5 minutes with room temperature PBS(+/+). Cells were then blocked in 10% animal serum of the host-species of the secondary antibody, diluted in 0.1% PBS(+/+)-Tween 20 (PBST) for 30 minutes. Primary antibody was diluted in 10% animal serum of the host-species of secondary antibody, diluted with 0.1% PBST overnight. Following primary incubation, cells were washed once for 5 minutes in PSB(+/+) before incubation in secondary antibody, diluted in 10% animal serum of the host-species of secondary antibody in 0.1% PBST for 1 hour. See supplementary Table 4 for primary and secondary antibody dilutions. Hoechst staining was completed where necessary (secondary only controls in the case of colocalization) for 5 minutes in PBS(+/+) followed two washes in PBS(+/+) for 5 minutes per wash. Cells were then mounted onto coverslips with Prolong Gold (P36934). Each experiment and their individual cell types included a secondary only control that was analysed with the same laser intensities as the treatment samples. Individual treatments were completed in three biological replicates. Primary and secondary antibodies and their concentrations are listed in Table 4.

**Table 4.**
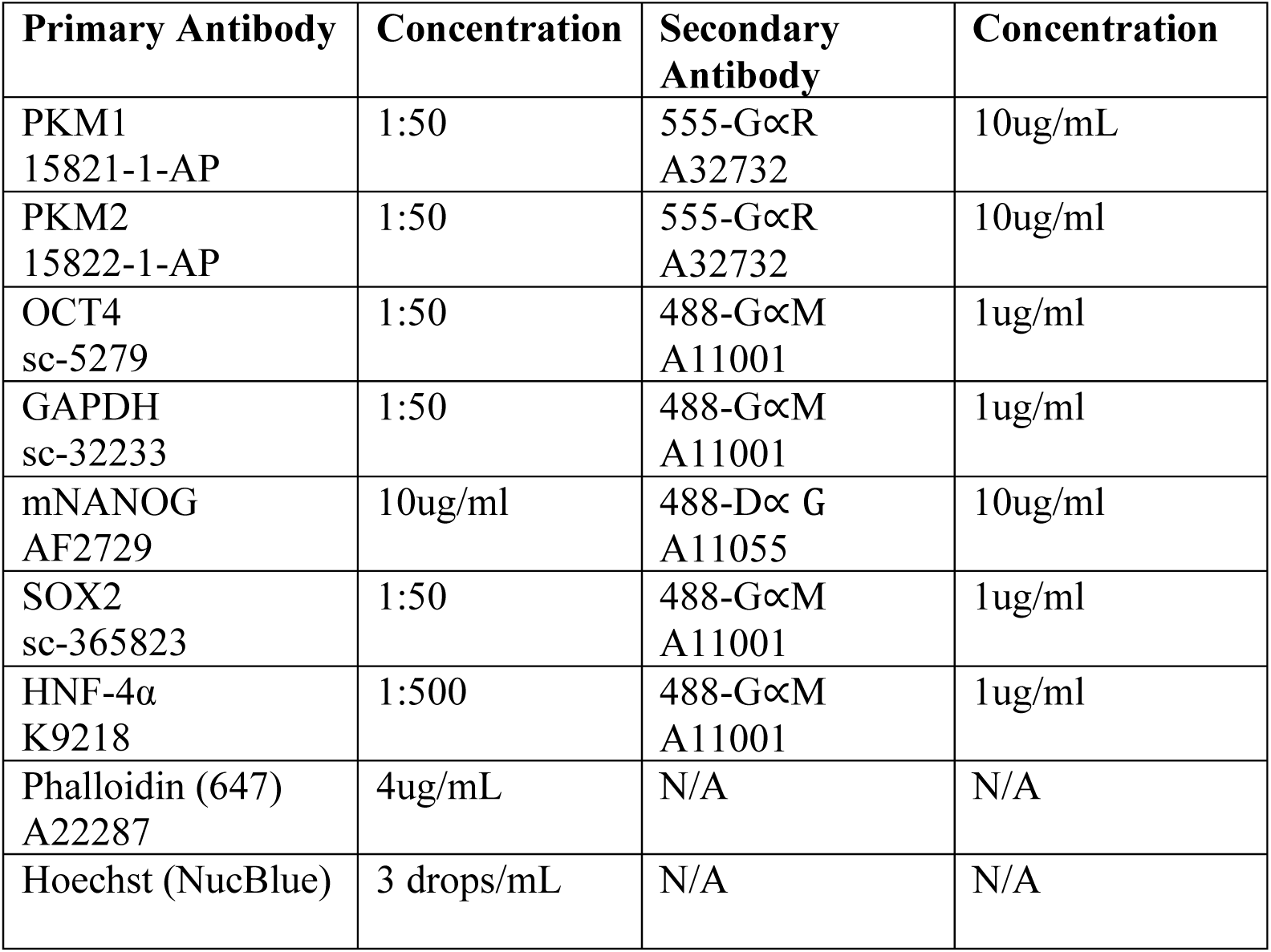
Immunofluorescence antibody and stain list.

| Primary Antibody | Concentration | Secondary Antibody | Concentration |
| --- | --- | --- | --- |
| PKM1<br>15821-1-AP | 1:50 | 555-GαR<br>A32732 | 10ug/mL |
| PKM2<br>15822-1-AP | 1:50 | 555-GαR<br>A32732 | 10ug/ml |
| OCT4<br>sc-5279 | 1:50 | 488-GαM<br>A11001 | 1ug/ml |
| GAPDH<br>sc-32233 | 1:50 | 488-GαM<br>A11001 | 1ug/ml |
| mNANOG<br>AF2729 | 10ug/ml | 488-Dα G<br>A11055 | 10ug/ml |
| SOX2<br>sc-365823 | 1:50 | 488-GαM<br>A11001 | 1ug/ml |
| HNF-4α<br>K9218 | 1:500 | 488-GαM<br>A11001 | 1ug/ml |
| Phalloidin (647)<br>A22287 | 4ug/mL | N/A | N/A |
| Hoechst (NucBlue) | 3 drops/mL | N/A | N/A |

### Colocalization: co-occurrence and correlation by immunofluorescence

Orthogonal projections of colony optimal slice generated image stacks were taken at 40x and 63x immersed in oil by a Zeiss LSM800 confocal microscope. Thresholds were set by optimized single stain samples (channel 488 - OCT4, GAPDH and channel 555 - PKM1, PKM2) exposed to all tested lasers and exposures. These exposures and laser intensities were tested against secondary antibody only controls. Double stains (PKM1/OCT4, PKM1/GAPDH, PKM2/OCT4 and PKM2/GAPDH) were taken in stacks containing full colonies and processed into orthogonal projections. The projections were then set to the predetermined co-localization thresholds (Costes thresholds were set when applicable and appropriate) as set from the single stain controls. Each treatment was analysed in at least biological triplicate and each biological replicate was examined for several technical replicates of different colonies within their respective samples. Double stained treatments were compared for co-occurrence and correlation using Manders’ Overlap Coefficient (MOC) and Pearson’s Correlation Coefficient (PCC) respectively. Additionally, we compared individual cells to whole colonies using airyscan processing under 63x magnification by confocal microscopy. This process increased the signal to noise ratio thus increasing signal resolution. PCC values were categorised within set ranges to a classification, that included: Correlation: very weak: -1 – -0.27, weak: -0.26 - 0.09, moderate: 0.1 – 0.48, strong: 0.49 – 0.84, and very strong: 0.85 – 1.0 (71). MOC values fall into set ranges of: Overlap: very weak: 0 – 0.49, weak: 0.50 – 0.70, moderate: 0.71 – 0.88, strong: 0.89 – 0.97, very strong: 0.98 – 1.0 (71). Statistical analysis included application of a two tailed Mann-Whitney test of mean ±SEM MOC and PCC scores run in at least three biological and technical replicates. Statistics of PCC and MOC treatments relative to the positive reference represent a two-way ANOVA with Sidak’s multiple comparisons test of mean±SEM PCC and MOC scores where ∝=0.05, n=3 biological replicates.

### Nuclear and cytoplasmic fractionation

Rapid isolation of nuclei from cells was completed using the REAP protocol (72). mESCs were grown to 90% confluency on 10 cm dishes. Prior to collection, culture medium was aspirated, and the cells were washed with ice-cold PBS(-/-) with 1X Protease Inhibitor Cocktail Set 1 (Calbiochem 539131). The PBS was aspirated, and the dish was placed on ice where 1 mL of PBS was added, and the cells were scraped and centrifuged for 10 seconds at 10,000 rpm. The supernatant was aspirated and resuspended in 900uL of ice-cold 0.1% Tergitol-NP-40 (Sigma NP-40S) in PBS(-/-) before being triturated 5 times. At this point a 300 μL total lysate sample was removed and stabilized in Laemmli buffer and vortexed. This sample was sonicated at 20 kHz for 2 pulses each 8 seconds long and the sample was then boiled for 1 minute and frozen prior to western blotting. The remaining NP-40 suspended sample was then centrifuged at 10,000 rpm for 10 seconds and 300 μL was removed as the cytoplasmic fraction. This fraction was stabilized in Laemmli buffer, vortexed and boiled for 1 minute before being frozen prior to western blotting. The remaining NP-40 suspended sample was aspirated and resuspended in 1mL 0.1%NP-40 in PBS(-/-) with 1X Protease Inhibitor Cocktail Set 1 before centrifuged at 10,000 rpm. The supernatant was discarded, and the pellet resuspended in water and Laemmli buffer before sonication at 20kHz for 2 pulses at 8 seconds per pulse. This nuclear fraction was boiled for 1 minute and frozen for future western blotting as described above. Antibody staining for control markers LAMIN A and ∝-TUBULIN and the markers of interest PKM1, pPKM2 and PKM2 were compared relative to total lane protein content by Ponceau staining (0.1% Ponceau, 5% acetic acid). Each cell type’s mean densitometry ± SEM was analyzed by applying a one-way ANOVA with Tukey’s multiple comparison test for unpaired, two tailed t-test respectively with three biological replicates.

## Supporting information

S1_raw_images

S1 Fig. mEpiLC generation and cell morphology.

S2 Fig. Secondary antibody only immunofluorescence controls for pluripotency markers.

S3 Fig. Secondary antibody only immunofluorescence controls for PKM1 and PKM2 and colocalization study.

S4 Fig. mESC PKM1, PKM2, OCT4 and GAPDH colocalization settings.

S5 Fig. mEpiLC PKM1, PKM2, OCT4 and GAPDH colocalization settings.

S6 Fig. mEpiSC PKM1, PKM2, OCT4 and GAPDH colocalization settings.

S7 Fig. mESC positive and negative colocalization controls.

S8 Fig. PKM1/2 colocalization within individual cells of mESC colonies.

S9 Fig. PKM1/2 colocalization within individual cells of mEpiLC colonies.

S10 Fig. PKM1/2 colocalization within individual cells of mEpiSC colonies

## Acknowledgments

This project was benefited through the training, advice and resources provided by Robarts’ London Regional Flow Cytometry Facility and Dr. Kristin Chadwick. Technical expertise and advisement from Courtney Brooks and Drs. Lin Zhao, Christie Vanderboor, Nicole Edwards, Ian Tobias, Jodi Garner, Amy Wong, and Dr. Cheryle Séguin aided to the progress of this work. Confocal and colocalization tips were graciously shared by Dr. Julia Abitbol. Pluripotent cell lines were gifted from Dr. Janet Rossant. This research was funded by a Canadian Institutes of Health Research operating grants to A.J.W. and D.H.B. and Natural Sciences and Engineering Research Council of Canada grant to D.H.B. The funders had no role in study design, data collection and analysis, decision to publish, or preparation of the manuscript.

## Supporting Information

### Supplementary Figures

**S1_raw_images**

Used and unused immunoblotting images. X denotes unused.

**S1 Fig. mEpiLC generation and cell morphology.** (A) Schematic depicting the generation of mEpiLCs from mESCs and the associated pluripotent states. (B) Phase contrast microscopy of mESCs, mEpiLCs (24, 48, 72, 96 and 120 hours) and mEpiSCs grown on MEFs. Images take using 10x Magnification and scale bars represent 250 and 300 *μm* (as indicated).

**S2 Fig. Secondary antibody only immunofluorescence controls for pluripotency markers.** Immunofluorescence of mESC, mEpiLC and mEpiSC stained for Hoechst, phalloidin and the secondary antibodies (Table 4) used throughout this study, assessed by confocal microscopy. Images taken using 40xmagnification and scale bars represent 20*μm*.

**S3 Fig. Secondary antibody only immunofluorescence controls for PKM1 and PKM2 and colocalization study.** Immunofluorescence microscopy of mESC, mEpiLC and mEpiSC stained for Hoechst, phalloidin and the secondary antibodies (Table 4) used throughout this study, assessed by confocal microscopy. Images taken using 40x magnification and scale bars represent 20*μm*.

**S4 Fig. mESC PKM1, PKM2, OCT4 and GAPDH colocalization settings.** Immunofluorescence microscopy of mESCs demonstrating single fluorescence images for PKM1, PKM2, OCT4 and GAPDH along with their respective thresholds. Images taken using 40x magnification and scale bars represent 20*μm*. Confocal laser channels labelled as 488 and 555 corresponding to treatments incubated with OCT4/GAPDH and PKM1/PKM2 respectively.

**S5 Fig. mEpiLC PKM1, PKM2, OCT4 and GAPDH colocalization settings.** Immunofluorescence microscopy of mEpiLCs demonstrating single fluorescence images for PKM1, PKM2, OCT4 and GAPDH along with their respective thresholds. Images taken using 40x magnification and scale bars represent 20*μm*. Confocal laser channels labelled as 488 and 555 corresponding to treatments incubated with OCT4/GAPDH and PKM1/PKM2 respectively.

**S6 Fig. mEpiSC PKM1, PKM2, OCT4 and GAPDH colocalization settings.** Immunofluorescence microscopy of mEpiSCs demonstrating single fluorescence images for PKM1, PKM2, OCT4 and GAPDH along with their respective thresholds. Images taken using 40x magnification and scale bars represent 20*μm*. White outlines represent area of analysis to exclude areas of MEF staining. Confocal laser channels labelled as 488 and 555 corresponding to treatments incubated with OCT4/GAPDH and PKM1/PKM2 respectively.

**S7 Fig. mESC positive and negative colocalization controls.** Immunofluorescence microscopy of mESCs demonstrating single and double stains for Hoechst, OCT4 and GAPDH along with their respective thresholds. Images taken using 40x magnification and scale bars represent 20*μm*. Confocal laser channels labelled as 405nm and 488nm corresponding to treatments incubated with Hoechst and OCT4/GAPDH respectively.

**S8 Fig. PKM1/2 colocalization within individual cells of mESC colonies.** Immunofluorescence microscopy of mESC colonies, colocalization analysis compared MOC and PCC of the total colony to that of a single cell. (A) PKM2 staining versus OCT4 and GAPDH in mESCs comparing orthogonal projections of whole colonies to individual cells by airyscan processing. (B) PKM1 staining versus OCT4 and GAPDH in mESCs comparing orthogonal projections of whole colonies to individual cells by airyscan processing. Images taken using 40x magnification with scale bars represent 20*μm* and 63x magnification with scale bars representing 5*μm*. Square boxes indicate areas of interest from the 40x for 63x magnification. White outlines around cells represents the area of analysis of the airyscanned images.

**S9 Fig. PKM1/2 colocalization within individual cells of mEpiLC colonies.** Immunofluorescence microscopy of mEpiLC colonies, colocalization analysis compared MOC and PCC of the total colony to that of a single cell. (A) PKM2 staining versus OCT4 and GAPDH in mEpiLCs comparing orthogonal projections of whole colonies to individual cells by airyscan processing. (B) PKM1 staining versus OCT4 and GAPDH in mEpiLCs comparing orthogonal projections of whole colonies to individual cells by airyscan processing. Images taken using 40x magnification with scale bars represent 20*μm* and 63x magnification with scale bars representing 5*μm*. Square boxes indicate areas of interest from the 40x for 63x magnification. White outlines around cells represents the area of analysis of the airyscanned images.

**S10 Fig. PKM1/2 colocalization within individual cells of mEpiSC colonies** Immunofluorescence microscopy of mEpiSC colonies, colocalization analysis compared MOC and PCC of the total colony to that of a single cell. (A) PKM2 staining versus OCT4 and GAPDH in mEpiSCs comparing orthogonal projections of whole colonies to individual cells by airyscan processing. (B) PKM1 staining versus OCT4 and GAPDH in mEpiSCs comparing orthogonal projections of whole colonies to individual cells by airyscan processing. Images taken using 40x magnification with scale bars represent 20*μm* and 63x magnification with scale bars representing 5*μm*. Square boxes indicate areas of interest from the 40x for 63x magnification. White outlines around cells represents the area of analysis of the airyscanned images.

## Notes

### Competing Interest Statement

The authors have declared no competing interest.

