## Supplementary figures and images for "Differential localization patterns of pyruvate kinase isoforms in murine naïve, formative and primed pluripotent states"

### S1 Fig. mEpiLC generation and cell morphology.

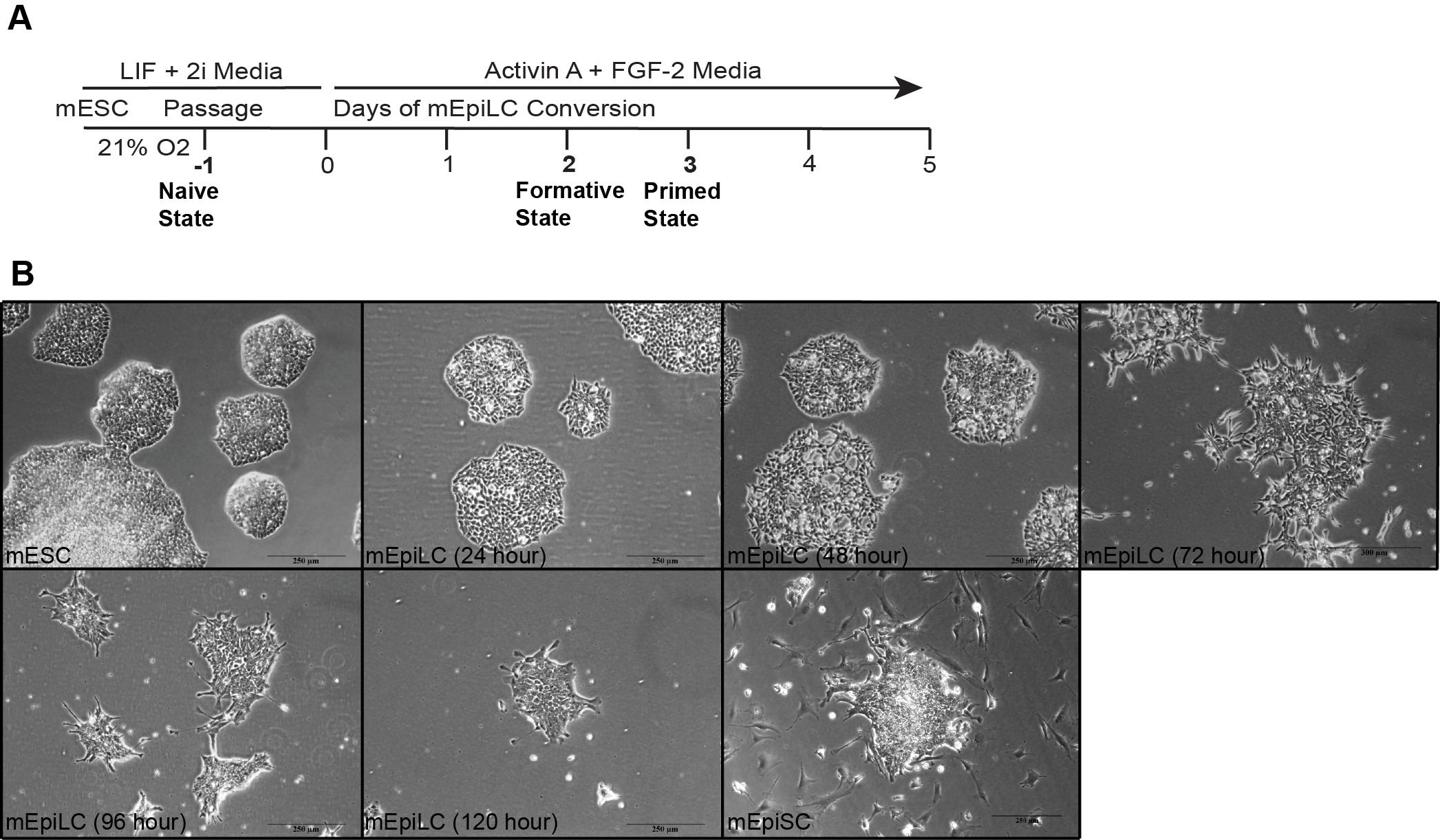

### S1_raw_images

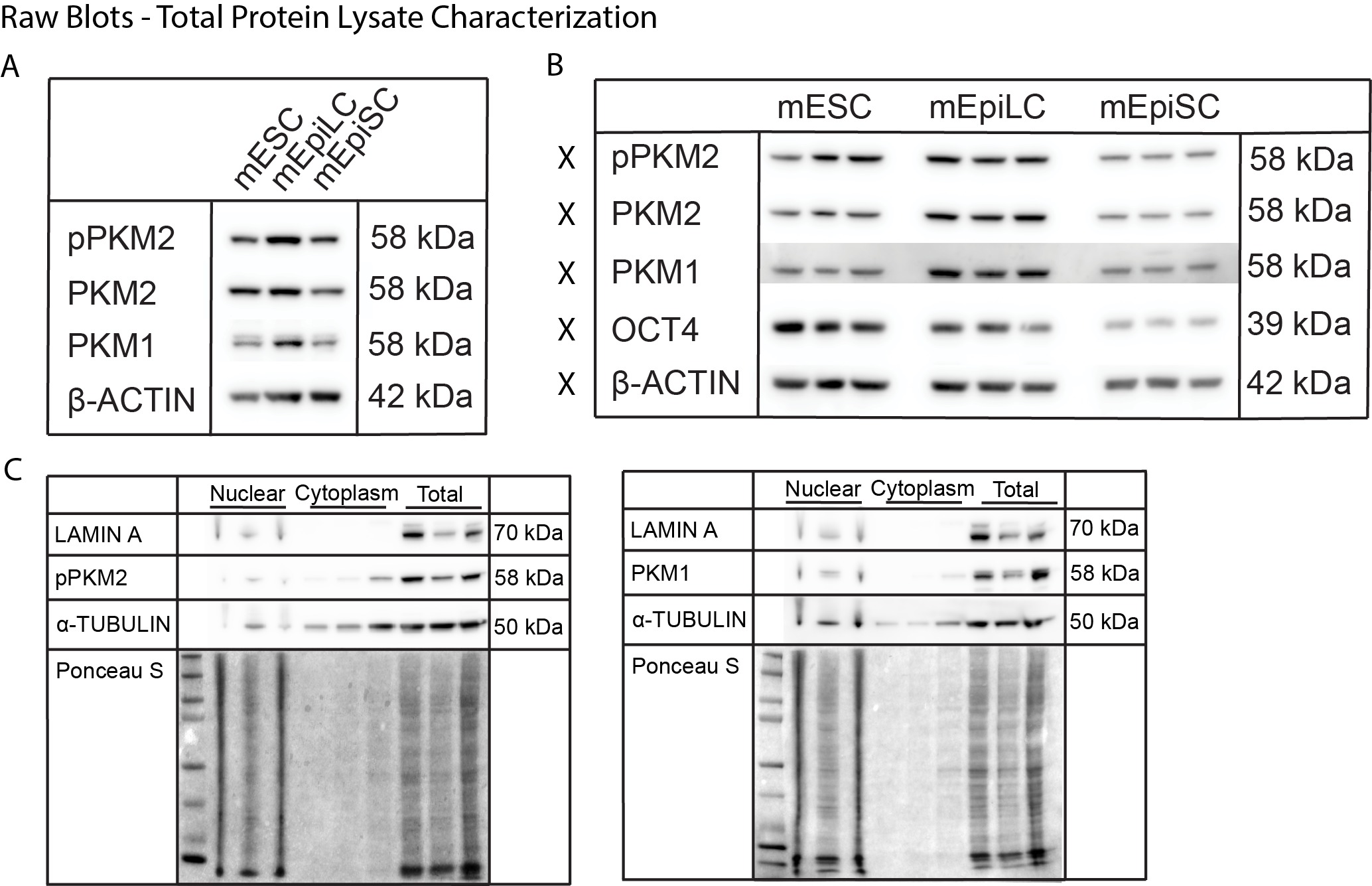

### S2 Fig. Secondary antibody only immunofluorescence controls for pluripotency markers.

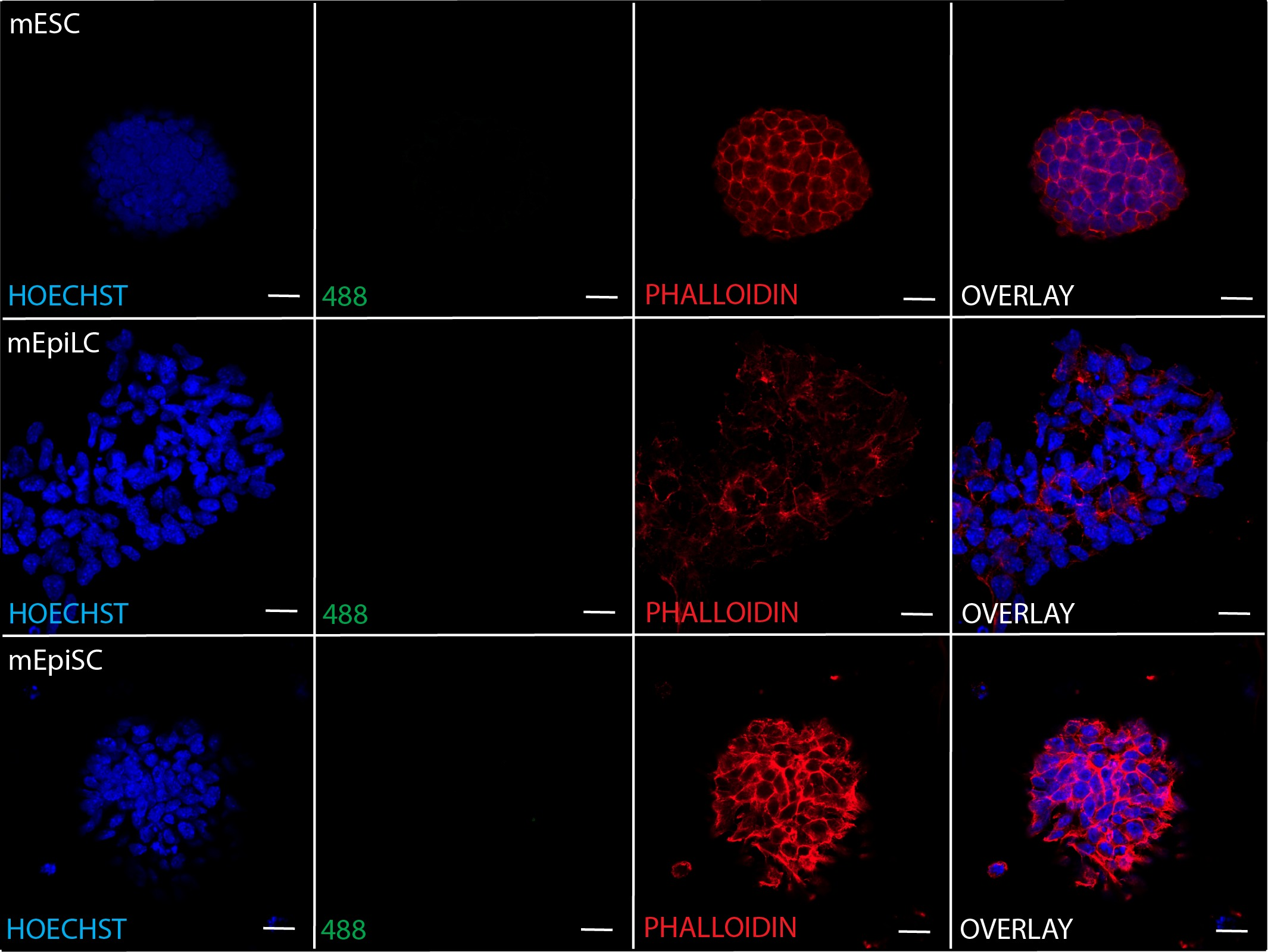

### S3 Fig. Secondary antibody only immunofluorescence controls for PKM1 and PKM2 and colocalization study.

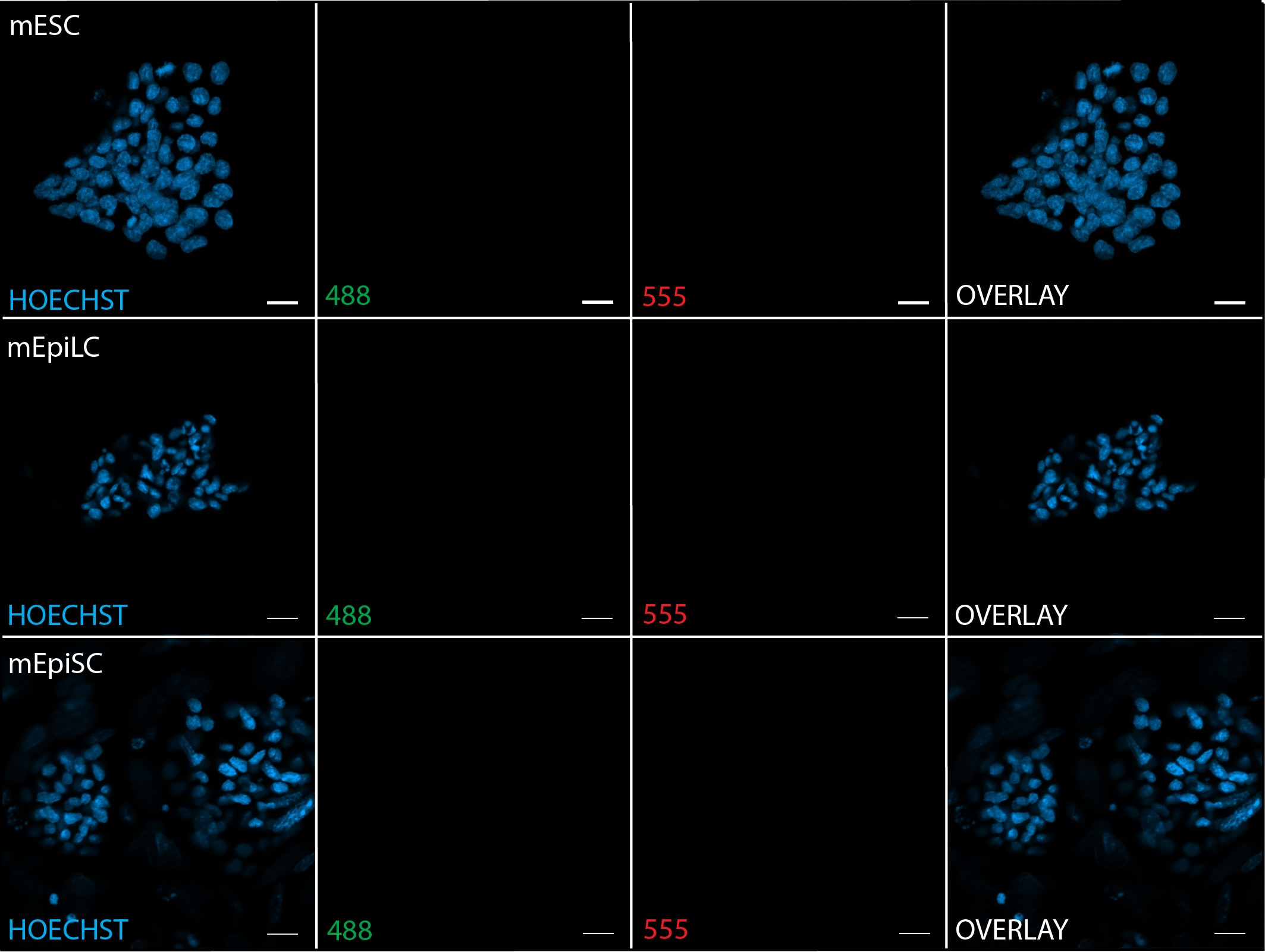

### S4 Fig. mESC PKM1, PKM2, OCT4 and GAPDH colocalization settings.

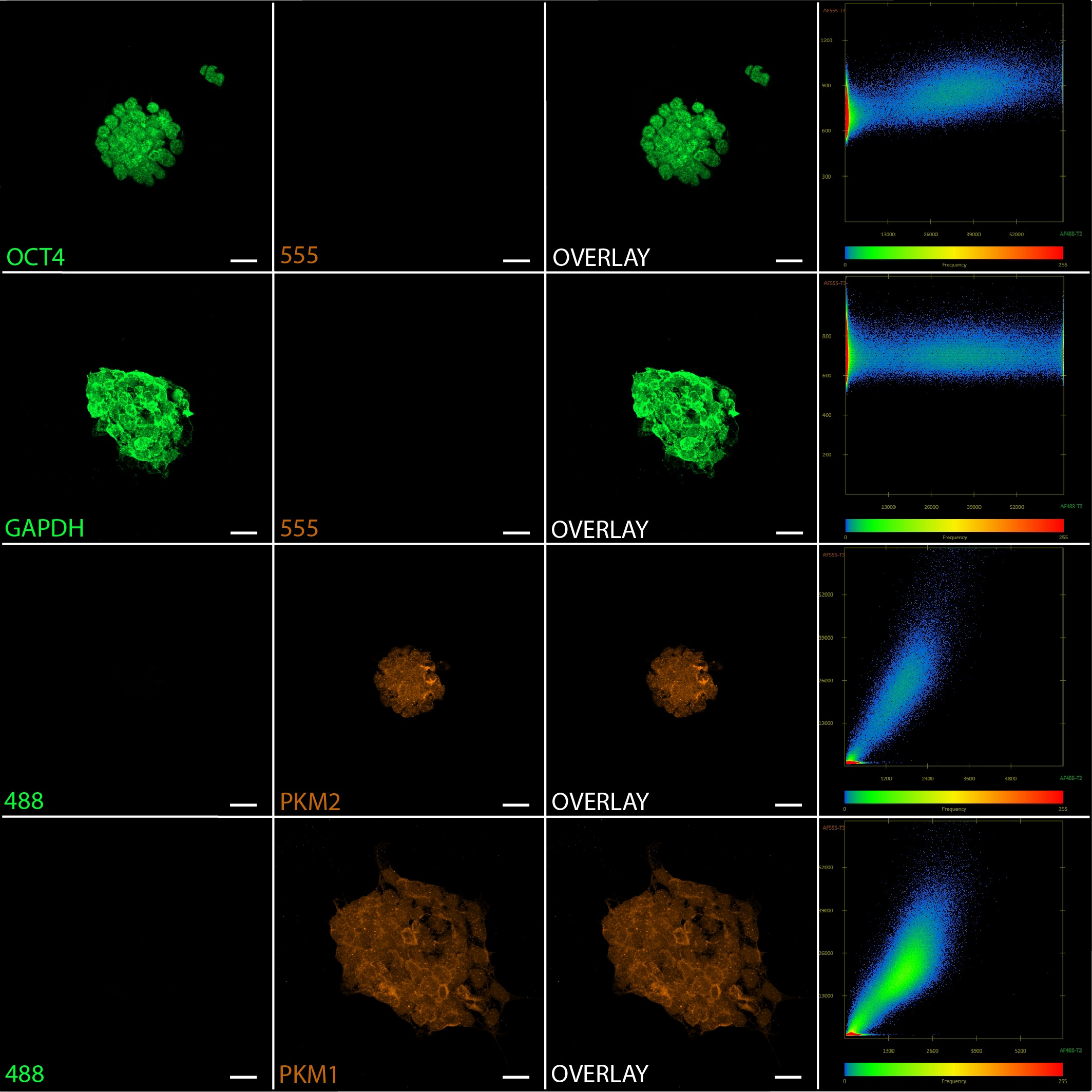

### S5 Fig. mEpiLC PKM1, PKM2, OCT4 and GAPDH colocalization settings.

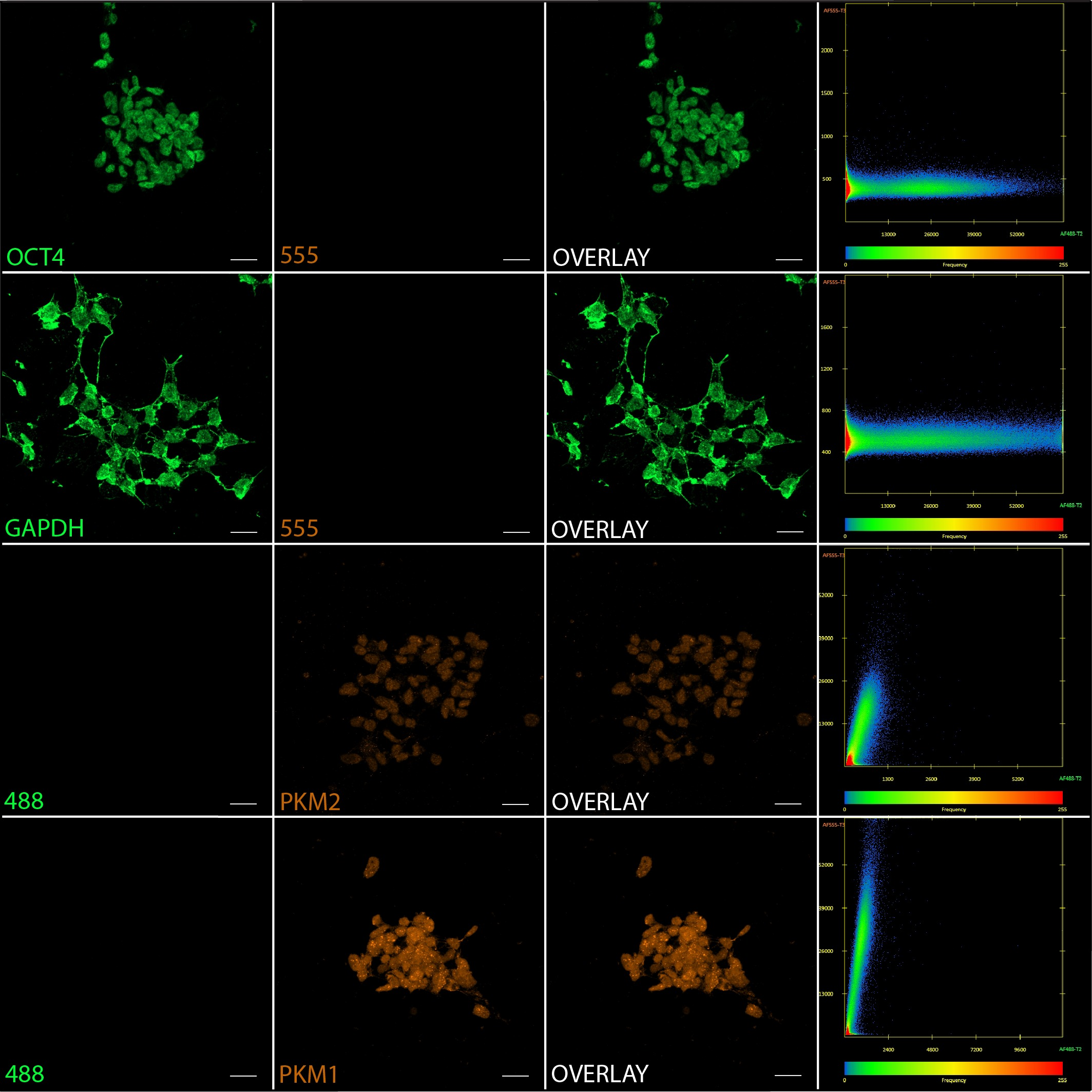

### S6 Fig. mEpiSC PKM1, PKM2, OCT4 and GAPDH colocalization settings.

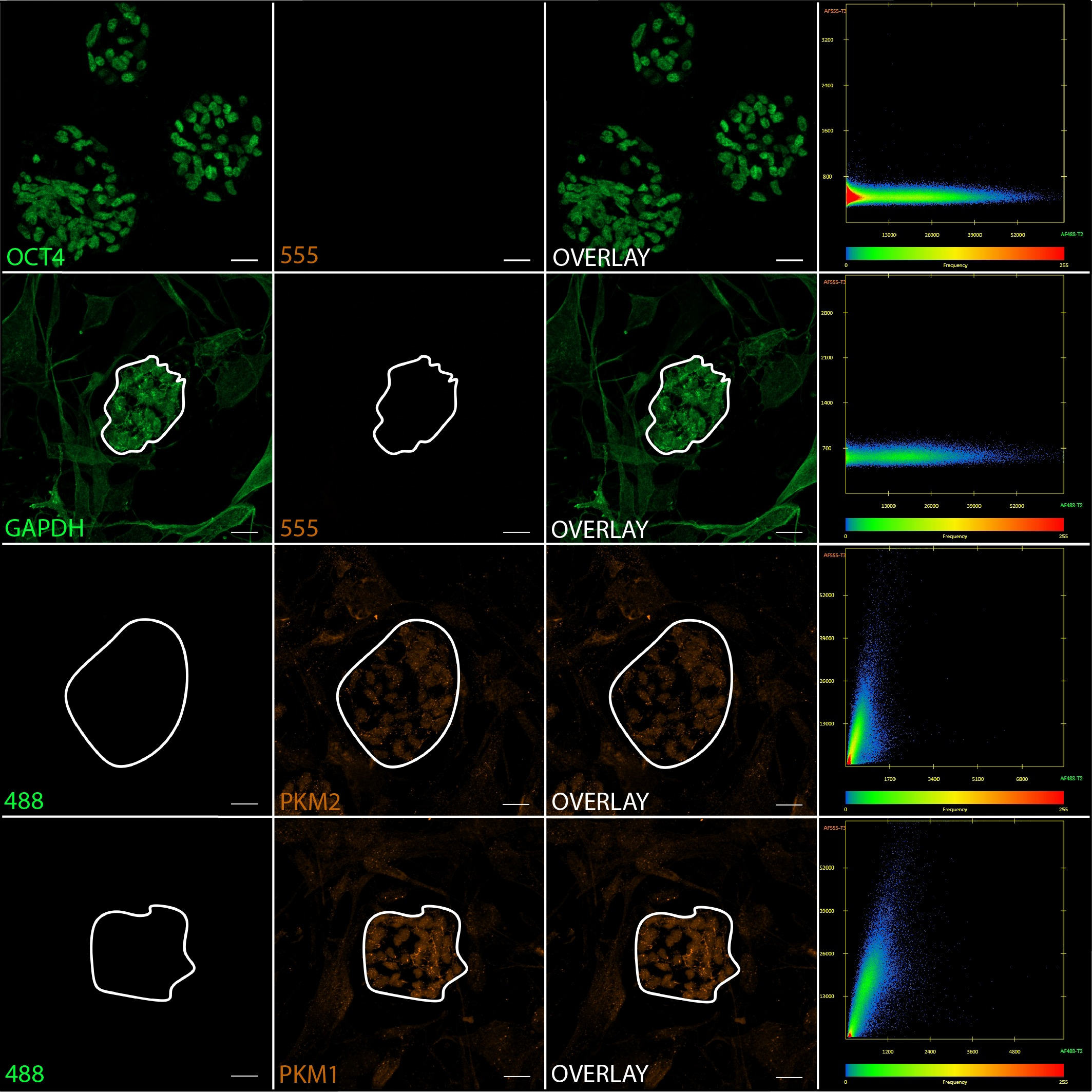

### S7 Fig. mESC positive and negative colocalization controls.

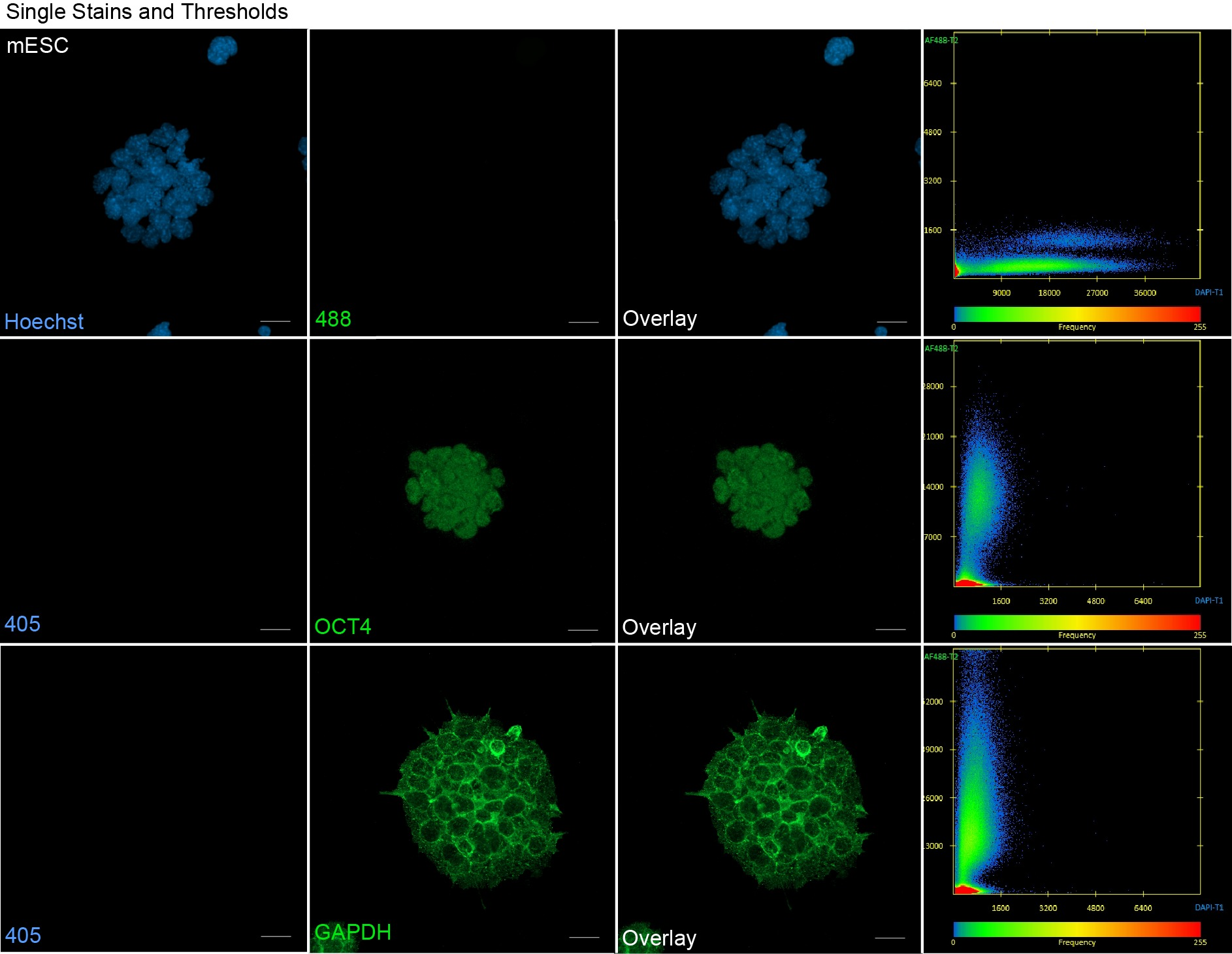

### S8 Fig. PKM1/2 colocalization within individual cells of mESC colonies.

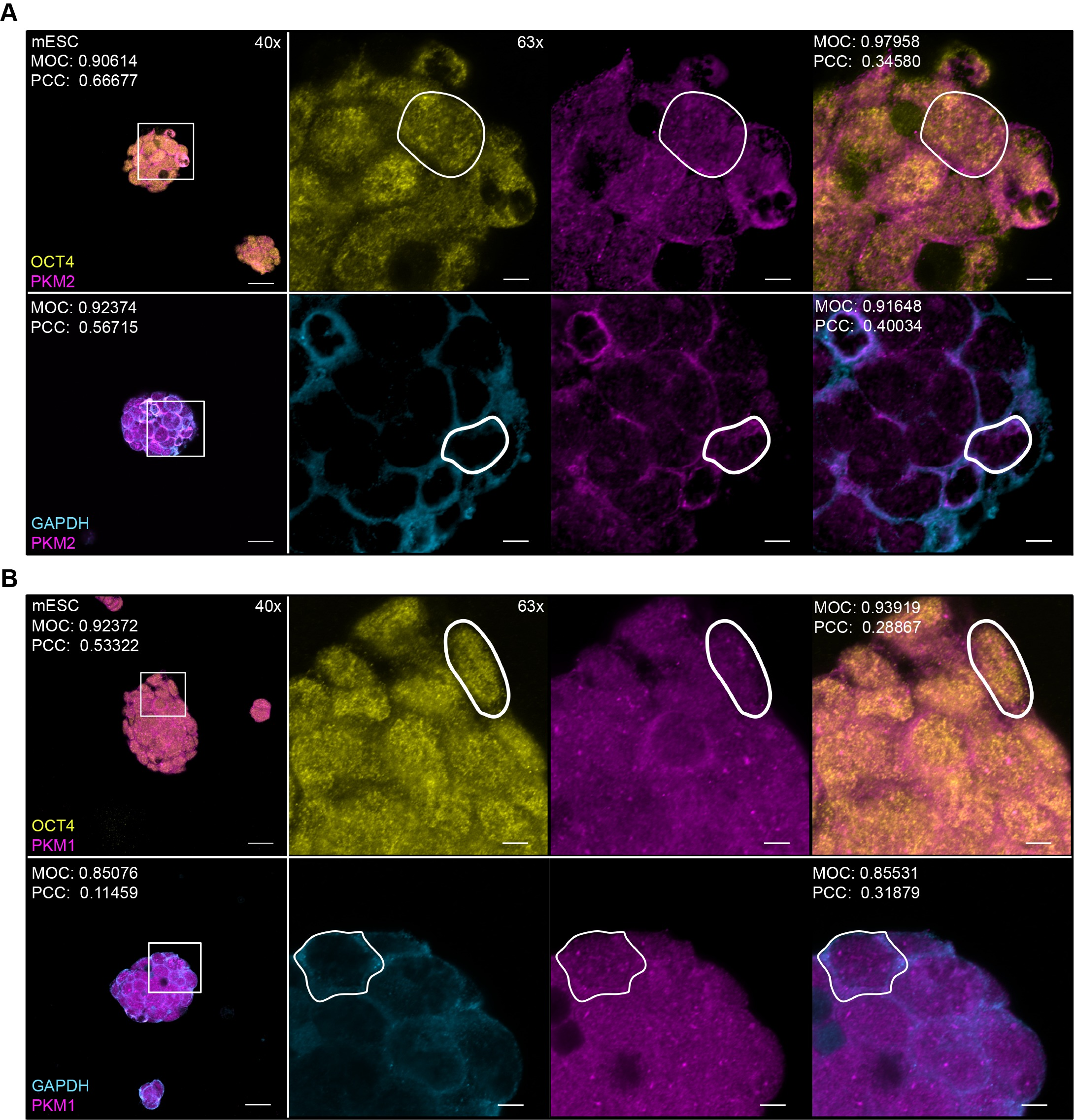

### S9 Fig. PKM1/2 colocalization within individual cells of mEpiLC colonies.

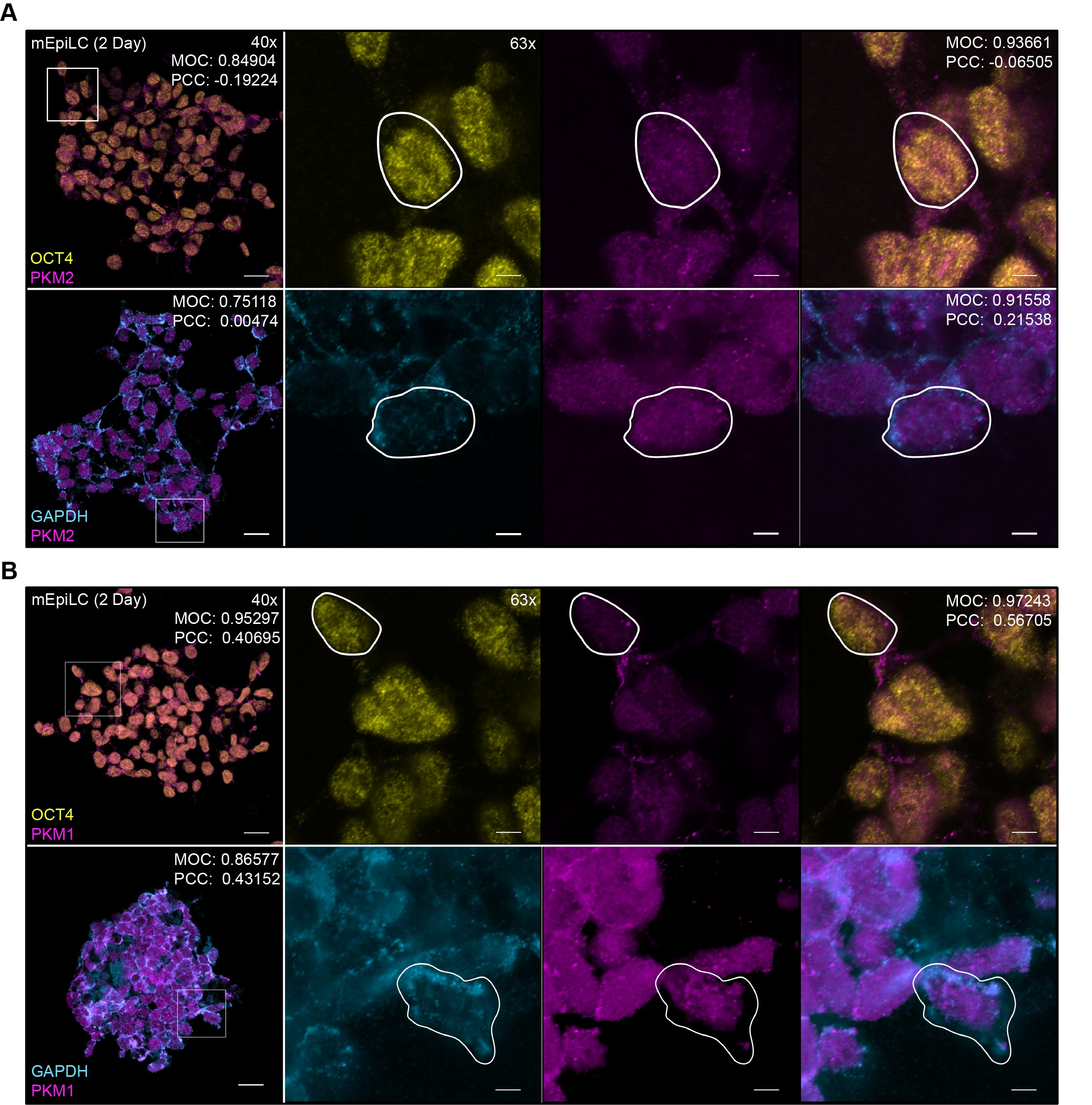

### S10 Fig. PKM1/2 colocalization within individual cells of mEpiSC colonies

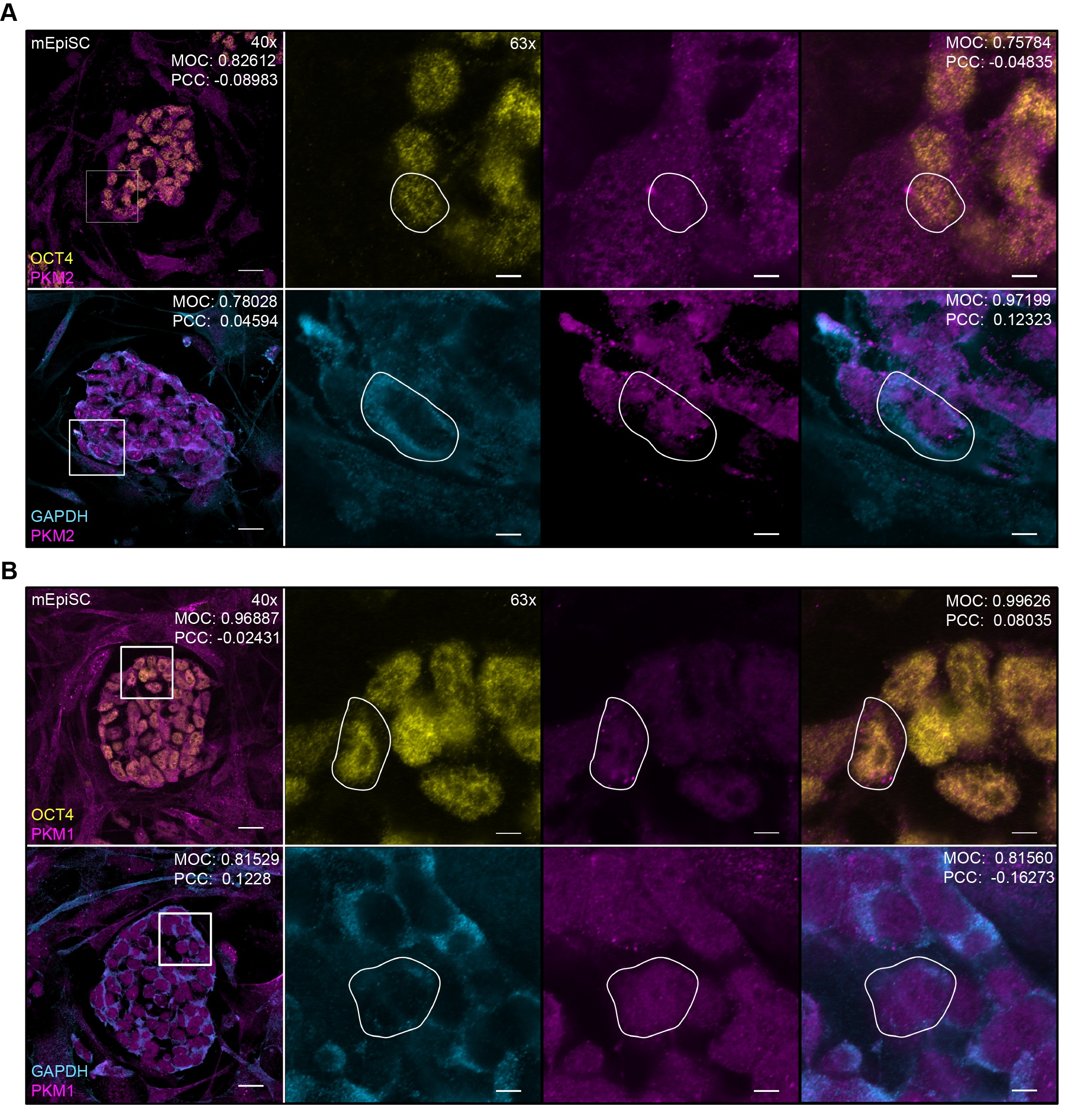
